# Forebrain projection neurons target functionally diverse respiratory control areas in the midbrain, pons and medulla oblongata

**DOI:** 10.1101/2020.08.21.260422

**Authors:** Pedro Trevizan-Baú, Rishi R. Dhingra, Werner I. Furuya, Davor Stanić, Stuart B. Mazzone, Mathias Dutschmann

**Affiliations:** The Florey Institute of Neuroscience and Mental Health, Discovery Neuroscience theme, The University of Melbourne, Parkville, VIC, 3010, Australia; Department of Anatomy and Neuroscience, The University of Melbourne, Parkville, VIC, 3010, Australia

**Keywords:** Pyramidal neurons, forebrain projection neurons, respiratory pattern generation, orofacial motor behaviors, volitional control of breathing, post-inspiration, delta, theta

## Abstract

Eupnea is generated by neural circuits located in the ponto-medullary brainstem, but can be modulated by higher brain inputs which contribute to volitional control of breathing and the expression of orofacial behaviors, such as vocalization, sniffing, coughing and swallowing. Surprisingly, the anatomical organization of descending inputs that connect the forebrain with the brainstem respiratory network remains poorly defined. We hypothesized that descending forebrain projections target multiple distributed respiratory control nuclei across the neuraxis. To test our hypothesis, we made discrete unilateral microinjections of the retrograde tracer Cholera toxin subunit B (CT-B) in the midbrain periaqueductal gray (PAG), the pontine Kölliker-Fuse nucleus (KFn), the medullary Bötzinger complex (BötC), pre-Bötzinger complex (pre-BötC) or caudal midline raphé nuclei. We quantified the regional distribution of retrogradely-labeled neurons in the forebrain 12-14 days post-injection. Overall, our data reveals that descending inputs from cortical areas predominantly target the PAG and KFn. Differential forebrain regions innervating the PAG (prefrontal, cingulate cortices, and lateral septum) and KFn (rhinal, piriform, and somatosensory cortices) imply that volitional motor commands for vocalization are specifically relayed via the PAG, while the KFn may receive commands to coordinate breathing with other orofacial behaviors (e.g. sniffing, swallowing). Additionally, we observed that the limbic or autonomic (interoceptive) systems are connected to broadly distributed downstream bulbar respiratory networks. Collectively, these data provide a neural substrate to explain how volitional, state-dependent, and emotional modulation of breathing is regulated by the forebrain.

## 1. Introduction

Studies on volitional control of breathing, which were mostly performed in humans, indicate that corticospinal pathways bypass the respiratory network of the brainstem and directly modulate respiratory motor pools in the spinal cord (Gandevia and Rothwell, 1987a,b; Corefiled et al., 1998; Butler, 2007; Pouget et al., 2018). In mammals, cortico-spinal pathways that specifically target spinal respiratory motor pools have been demonstrated (Rikard-Bell et al., 1985). However, descending anatomical pathways that target nuclei of the primary respiratory rhythm and pattern generating network remain poorly defined despite functional evidence that activity within the respiratory network is modulated by behavioral commands (Orem and Netick, 1986; Chang, 1992). Cognitive and behavioral motor commands modulate respiratory activities during a variety of volitional orofacial behaviors including vocalization (speech), swallowing, chewing, coughing, sneezing, sighing, and sniffing (Davis et al., 1996; Nanoka et al., 1999; Jean 2001; Ludlow, 2005; Sherwood et al., 2005; Deschênes et al., 2012; Moore et al., 2013; Moore et al., 2014; Laplagne, 2018; McElvain et al., 2018). Many of these orofacial behaviors depend on the recruitment of upper airway muscles that regulate airway patency during inspiration and expiration (Dutschmann and Paton, 2002). Control of expiratory airflow during the post-inspiratory phase of respiration is essential for the mediation of vocalization, swallowing and expulsive expiratory behaviour (Dutschmann et al., 2014). Since the primary motor networks that control post-inspiration are distributed in the brainstem (Dhingra et al., 2019a,b; Dhingra et al., 2020), we hypothesized that crucial descending forebrain projections target respiratory control nuclei above the spinal cord. In addition, we assumed that orofacial behaviors that involve augmented inspiratory activity (e.g. sighing, sniffing) may also recruit brainstem nuclei that generate and/or modulate the inspiratory rhythm, instead of overwriting ongoing repiratory activity with cortico-spinal motor commands.

To test this working hypothesis, we used the retrograde tracer cholera toxin subunit B (CT-B) to characterize the topography of descending monosynaptic projections from the forebrain to five anatomically distinct respiratory areas in the midbrain and ponto-medullary brainstem that have established function in the modulation or generation of respiratory activity. The descending forebrain connectivity of the periaqueductal gray (PAG) was analyzed because it has a profound role in the modulation of respiration during vocalization and defensive behaviors (Zhang et al., 1994; Subramanian et al., 2008a,b; Subramanian and Holstege, 2014; Dampney, 2015; Faul et al., 2019), but has no breath-by-breath role in respiratory rhythm and pattern generation (Farmer et al., 2014). Because the PAG has established connectivity with various forebrain nuclei (Dampney et al., 2013), it also serves as an important control for the analysis of the additional respiratory brain areas studied. The pontine Kölliker-Fuse nucleus (KFn) was investigated because it regulates the inspiratory-expiratory phase transition and is involved in the breath-by-breath formation of the respiratory motor pattern (Caille et al., 1981; Wang et al., 1993; Dutschmann and Herbert, 2006; Smith et al., 2007; Mörschel and Dutschmann, 2009). Moreover, the KFn has major implications in the control of laryngeal adductor function during breathing (Dutschmann and Herbert 2006) and orofacial behaviors (Dutschmann and Dick, 2012). The pre-Bötzinger complex (pre-BötC) was chosen for its essential role in inspiratory rhythm generation (Smith et al., 1991; Feldman and Del Negro, 2006; Del Negro et al., 2018) and the neighbouring Bötzinger complex (BötC) was targeted because of its proposed function as an essential part of respiratory rhythm generating circuit (Burke et al., 2010; Smith et al., 2013; Marchenko et al., 2016). Finally, descending forebrain inputs to nuclei of the caudal raphé were analyzed since these serotonergic neurons have important neuromodulatory action on the respiratory motor pattern (Holtmann et al., 1986, Lindsey et al., 1998; Richter et al., 2003, Besnard et al., 2009; Hodges and Richerson; 2010).

## 2. Meterials and methods

### 2.1. Animals

Adult Sprague-Dawley rats of either sex (*n*=28, weight range: 280-350g) were used for this study. All animals were housed under a 12:12 h light/dark cycle, with free access to lab chow (Ridley Corporation Limited, Australia) and water. Experiments followed protocols approved by the Florey Institute of Neuroscience and Mental Health Animal Ethics Committee and performed in accordance with the National Health and Medical Research Council of Australia code of practice for the use of animals for scientific purposes.

### 2.2. Surgery

For tracer microinjections, rats were initially anaesthetized with isoflurane (5% v/v in oxygen). After mounting in a stereotaxic apparatus (TSE systems, Bad Homburg, Germany), anaesthesia was maintained with isoflurane (∼2% in oxygen) via a nose cone. After the rats were placed with the skull in a flat position in the stereotaxic frame, a craniotomy was performed. Surgeries were performed under an aseptic technique (iodine antiseptic solution). A midline incision exposed the skull between bregma and the interaural line, and a small burr hole was drilled according to coordinates defined relative to bregma (Table 1). Using a 1 µl Hamilton syringe (25s-gauge needle), 150nL of 1% CT-B (1 mg/mL; Invitrogen, OR, USA) was pressure injected unilaterally in the following brain nuclei: PAG (dorsolateral and ventrolateral columns; *n*=5, plus 3 near-miss injections), KFn (*n*=5, plus 2 near-miss injections), BötC (*n*=3, plus 1 near-miss injection), pre-BötC (*n*=3, plus 1 near-miss injection) and caudal raphé nuclei (raphé pallidus [RPa], raphé magnus [RMg], and raphé obscurus [ROb]; *n*=3, plus 2 near-miss injections). The total volume was injected at a rate of 20 nL/min, and CT-B injections were made on the left side of the brain. To avoid bleeding after injection through the transverse sinus for medullary targets (for BötC, pre-BötC, and caudal raphe injections), the syringe was angled and inserted from a more rostral location to reach the specific coordinates (for details, see Table 1). After the injection, the syringe remained in the brain tissue for at least 10 min and was withdrawn at 1mm/min to minimize non-specific spread of the tracer in brain tissue along the injection tract (Finkelstein et al, 2000). Immediately after surgery, animals received 3mg/kg of the anti-inflamatory drug Meloxicam (Tory-Ilium, NSW, Australia; 5mg/mL, deliveried subcutaneously). The animals were allowed to recover for 12-14 days before transcardial perfusion.

**Table 1.**
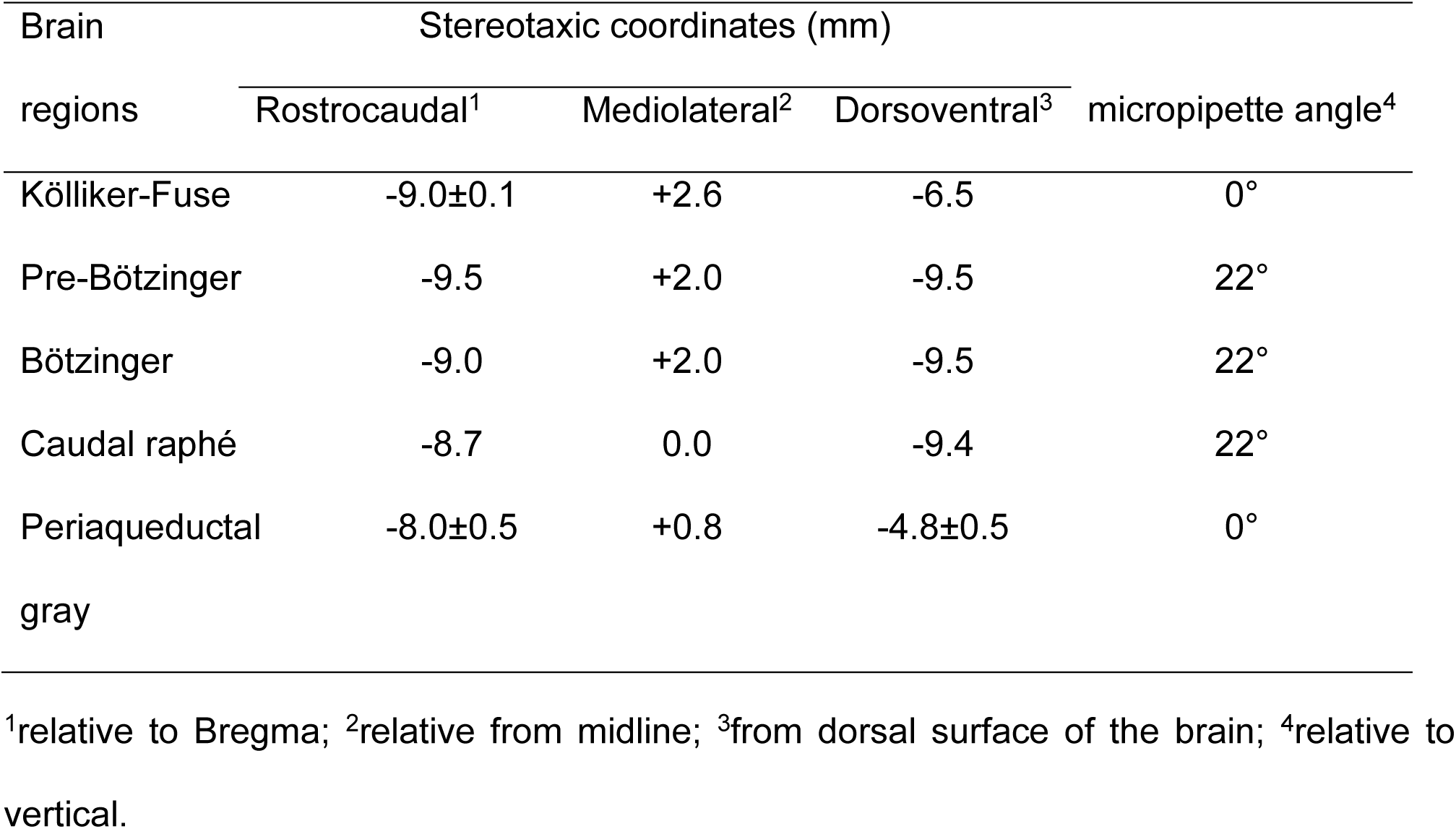
Stereotaxic coordinates for CT-B microinjections.

### 2.3. Tissue preparation and immunohistochemistry

Twelve to fourteen days after injection, the rats were deeply anaesthetized with sodium pentobarbitone (Ilium Pentobarbitone, Troy Laboratories, Smithfield, NSW, Australia, 100 mg/kg i.p.) and transcardially perfused with 60 mL Ca^2+^-free Tyrode’s buffer (37 °C), followed by 60 mL 4% paraformaldehyde (PFA, Sigma-Aldrich) containing 0.2% picric acid (Sigma) diluted in 0.16 M phosphate buffer (Merck KGaA, Darmstadt, Germany; pH 7.2, 37 °C) and finally, an additional 300 mL PFA/picric acid solution at 4°C. Brains were removed from the skull and post-fixed in PFA/picric acid solution for 90 min at 4°C. Next, the brains were immersed for 72 h in 0.1 M phosphate buffer (pH 7.4) containing 15% sucrose, 0.01% sodium azide (Sigma) and 0.02% bacitracin (Sigma) for cryprotection. A small incision from the olfactory bulbs to the brainstem was made in the hemisphere contralateral to the CT-B injection site to allow for subsequent measurements of the ipsilateral and contralateral distribution of projection neurons (Fig. 2). After postfixation and cryoprotection, brains were rapidly frozen using liquid carbon dioxide. Finally, brains we stored at -80 °C until cryo-sectioning.

**Figure 1.**
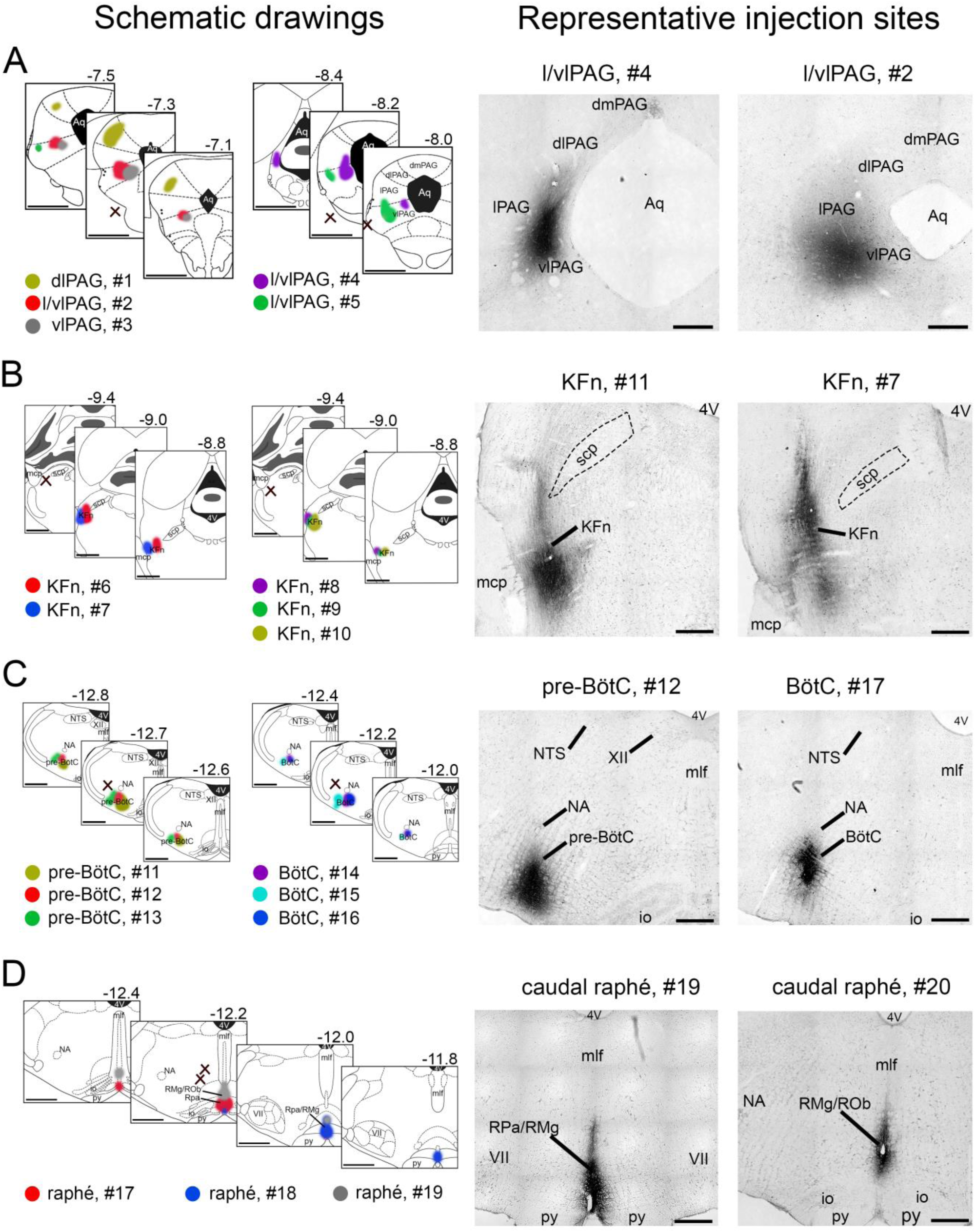
Schematic drawings (left) and photomicraphs (right) illustrate the anatomical location of CT-B microinjections in functionally diverse respiratory control areas: the periaqueductal gray **(A)**, the Kölliker-Fuse nucleus **(B)**, the pre-Bötzinger and Bötzinger complexes **(C)**, and the caudal raphé nuclei **(D)**. Left: schematic drawings depict the location and dimensions of all CT-B injections in the brainstem target areas relative to bregma. In this and in Figures 2, 5, 7, 9, 10, & 12, colors code specific on-target injections; and black x’s indicate near-miss, off-target injections (see results). Right: photomicraphs of representative injection sites. *Abbreviations*: 4V = fourth ventricle; BötC = Bötzinger complex; KFn = Kölliker-Fuse nucleus; mcp = middle cerebellar peduncle; io = inferior olive; mlf = medial longitudinal fasciculus; NA = nucleus ambiguus; NTS = solitary tract nucleus; PAG = periaqueductal gray; pre-BötC = pre-Bötzinger complex; py = pyramidal tract; RMg = raphé magnus; Rob = raphé obscurus; RPa = raphé pallidus; scp = superior cerebellar peduncle; XII = facial motor nucleus. *Scale bars*: 1 mm (schematic drawings on the left); 200 µm (representative photographs on the right).

**Figure 2.**
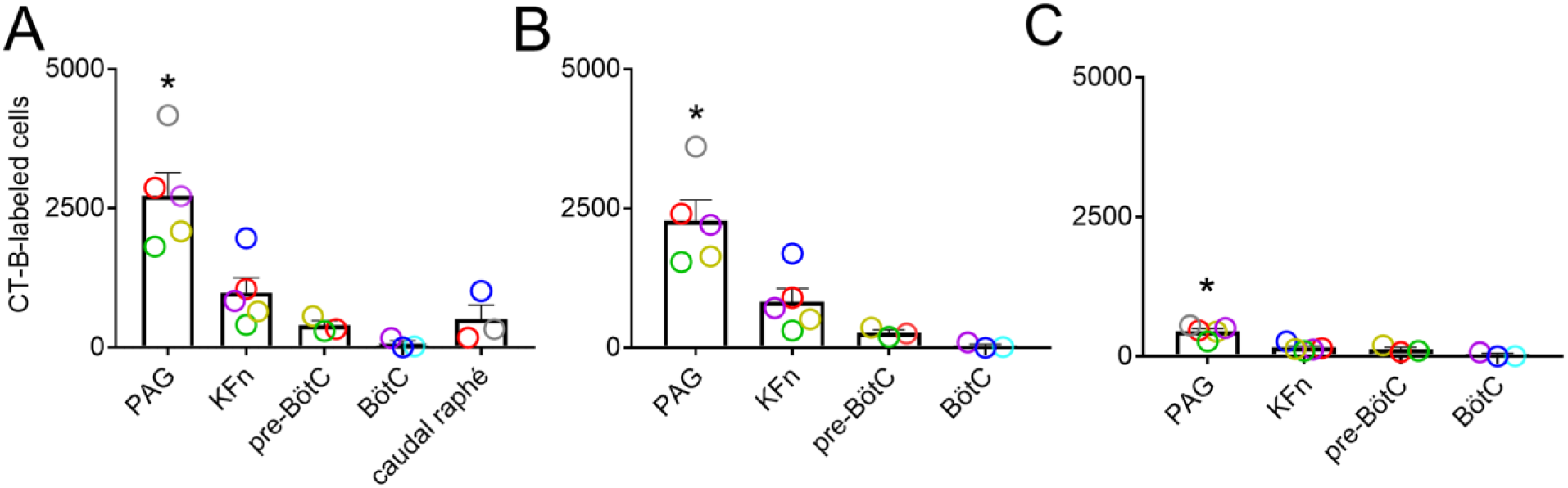
Bar graphs of the total **(A)**, ipsilateral **(B)** and contralateral **(C)** numbers of retrogradely CT-B-labeled neurons in the forebrain after microinjections in PAG, KFn, pre-BötC, BötC, and caudal raphé nuclei. In A, B, & C color-coded circles (experimental cases) align with those Figure 1; and zero laterality for caudal raphé nuclei. All values are expressed as mean ± standard error of the mean. The greatest number of CT-B retrogradely labeled forebrain neurons were detected from injections in the PAG compared to those in the KFn, pre-BötC, BötC, and caudal raphé nuclei (One-way ANOVA followed by Tukey’s multiple comparison test, *p < 0.05). *Abbreviations*: BötC = Bötzinger complex; KFn = Kölliker-Fuse nucleus; PAG = periaqueductal gray; pre-BötC = pre-Bötzinger complex.

Serial coronal sections (40 µm thickness) of the entire brain (from the olfactory bulb to the spino-medullary junction) were cut using a cryostat (Leica CM1850, Leica Microsystems), and stored in a cryo-protectant solution [30% v/v ethyleneglycol (Merck); 15% w/v sucrose; 35% v/v 0.1M phosphate buffer; 35% v/v distilled H2O] at −20°C. Brain sections were cut serially, and sections processed for immunohistochemistry were separated by 120 µm. Freely floating sections were washed in 0.01 M PBS, followed by incubation in hydrogen peroxide for 20 min to block endogenous peroxidase activity. Next, sections were incubated for 24 h at room temperature (RT) with a goat anti-CT-B antibody (List Biological Laboratories, CA, USA; catalogue no #112; 1:10,000) diluted in PBS containing 0.3% Triton X-100 and 0.5% BSA (Sigma). The goat anti-CT-B primary antibody recognizes the B-subunit of cholera toxin. Sections were then washed in 0.01 M PBS and blocked with 5% normal donkey serum (NDS) in 0.01M PBS for 1 hr at RT. Sections were immediately incubated in the corresponding secondary antibody (anti-sheep biotin, Jackson ImmunoResearch Laboratories, West Grove, PA; 1:500 in 0.01 M PBS) for 1h at RT. Next, sections were washed in 0.01 M PBS and incubated in an ABC kit (Vectastain® Elite ABC-HRP Kit) for 1h at RT. Finally, sections were washed in 0.01 M PBS and subsequently incubated in diaminobenzidine (DAB) substrate (1:10, Roche Diagnostics Mannheim, Germany) for 4 min, followed by an incubation in 1% H_2_O_2_ for 4 min (Stanic et al, 2003). Sections were then washed 3x in 0.01 M PBS and mounted on slides coated with 0.5% gelatin (Sigma) and 0.05% chromium (III) potassium sulphate dodecahydrate (Merck), and left to dry overnight. Slides were coverslipped using DPX (Sigma-Aldrich).

### 2.4. Data analysis and image processing

We used a brightfield microscope (Leica Biosystems) to identify and document the location of the midbrain and brainstem injection sites. Injection sites were photographed and plotted on semi-schematic drawings of the respective sections containing the PAG, KFn, BötC, pre-BötC and caudal raphé nuclei (Fig. 1). We used the following criteria to classify injections of the adjacent BötC and pre-BötC nuclei in the medulla. The center of the BötC injection was located 0.0-0.5mm to caudal edge of the facial motor nucleus (VII) where the medial longitudinal fasciculus (mlf) in the midline is ventral-dorsally prolonged till the fourth ventricle (4V); whereas the center of the pre-BötC injection sites was located 0.5-0.8mm to caudal edge of the facial motor nucleus where the mlf is less ventral-dorsally prolonged in the midline. Another landmark for the pre-BötC location was the presence of the hypoglossal nucleus (XII) in the dorsomedial region of the section, which is not present in the BötC sections (Fig. 1C).

Representative images were taken using a digital camera (Leica DFC7000T) mounted to the microscope (Leica DM6B LED). Adobe Photoshop CC19 software (Adobe Systems Inc., San Jose, CA) was used to assemble representative images, and to optimize brightness and contrast of the digital images to best represent sections viewed under the microscope.

Retrogradely labeled neurons in specific forebrain areas were quantified by counting the number of CT-B labeled neuronal cell bodies on sections separated by 120 µm across the neuraxis for all experimental cases. Near-miss injections served as controls. CT-B labeled cell bodies were recognized by the presence of black punctate granules in the neuronal cell bodies (somata) and processes, and the absence of immunoreactive cell nuclei. The background staining of neurons was characterized by light and diffuse DAB staining of somata including the cell nuclei. The specific location and distribution of retrogradely labeled forebrain projection neurons in cortical and sub-cortical areas are classified according to the rat brain atlas of Paxinos and Watson (2007).

We used a one-way analysis of variance (GraphPad Prism, version 7.02; GraphPad Software; San Diego, CA) followed by Tukey’s multiple comparison test to determine the statistical significance of the total, ipsilateral and contralateral numbers of CT-B-labeled neurons between the brainstem target areas (i.e. PAG, KFn, pre-BötC, BötC and caudal raphé nuclei; Fig. 2). Values are given as mean ± standard error of the mean (SEM) and *p*-values less than 0.05 were considered statistically significant.

## 3. Results

### 3.1. Injection sites

Microinjections of the retrograde tracer CT-B were anatomically confined to discrete injection sites in all target areas (see representative photographs, Fig. 1) or around the target areas (‘near-miss injections’). The latter are illustrated as black crosses in the semi-schematic drawings (Fig. 1, *left panel*).

CT-B injection sites in columns of the midbrain PAG were confined to the vlPAG (n=4) and the dlPAG (n=1; Fig. 1A). The rostrocaudal levels of the injections were identified by the size and location of the aqueduct (Bandler et al.,1991; Carrive, 1993). Three injection sites were centered in the rostral PAG (cases #1-3), whereas the other two cases (#4, 5) were located more caudally. We also report three near-miss injections, which were localized ventrolaterally to the PAG (Fig. 1A, *black x’s*).

CT-B injections in the pontine KFn, located ventral to the lateral tip of the superior cerebellar peduncle (scp), were restricted to the rostral and intermediate regions of the KFn (cases #6-10, Fig. 1B), and strongly overlap with the core circuitry of pontine respiratory group in rodents (Dutschmann and Herbert 2006). Two near-miss injections were found slightly ventrocaudally to the pontine KFn (Fig. 1B, *black x’s*).

Anatomical locations of the medullary pre-BötC (cases #11-13) and BötC (cases #14-16, Fig. 1C) were localized ventral to the nucleus ambiguus (NA) and caudal to the facial nucleus (VII) in the medulla oblongata. Two near-miss injections were located dorsolaterally to the NA (Fig. 1C, *black x’s*).

Injection sites for cases #17-19 were confined to the caudal midline raphé (Fig. 1D), which is sub-divided in three regions: raphé obscurus (ROb), pallidus (RPa) and magnus (RMg). Injections in the caudal raphé were located rostral to the BötC/pre-BötC brainstem sections. Case #17 was predominantly localized to the RPa. Case #18 was more rostral to case #17 and localized to the RPa/RMg. Case #19 was localized to the RMg/ROb. Two near-miss injections were located lateral to the caudal midline raphé (Fig. 1D, *black x’s*).

### 3.2. Cumulative distribution and laterality of descending forebrain projections

Quantitative analysis of the total numbers of CT-B-labeled neurons following injections in the midbrain PAG, pontine KFn, medullary pre-BötC, BötC, or the caudal raphé revealed that the highest number of CT-B-labeled cells in the forebrain projected monosynaptically to the PAG (p < 0.05; Fig. 2). Injections in the pontine KFn resulted in the second highest number of CT-B labeled neurons in the forebrain, followed by the medullary pre-BötC, and the caudal raphé. The lowest numbers of CT-B-labeled neurons were observed after injections in the medullary BötC.

Analysis of the ipsilateral and contralateral distribution of the CT-B-labeled neurons demonstrates that the majority of the CT-B-labeled neurons after PAG and KFn injections were located in the ipsilateral hemisphere (PAG, 83.1 ± 3.1%; KFn, 82.5 ± 4.6%). For injections in the pre-BötC and BötC, we respectively observed 69.8 ± 6.5% and 60.7 ± 6.0% of CT-B-labeled neurons were located ipsilaterally.

CT-B-labeled cell bodies in cortical regions were almost exclusively found in the layer V/VI, and displayed the characteristic morphology of cortical pyramidal neurons (Fig. 3). It is also important to note that CT-B-labeled neurons in the hippocampus, olfactory bulb, basal ganglia and cerebellum were not detected in any experiments reported in this study.

**Figure 3.**
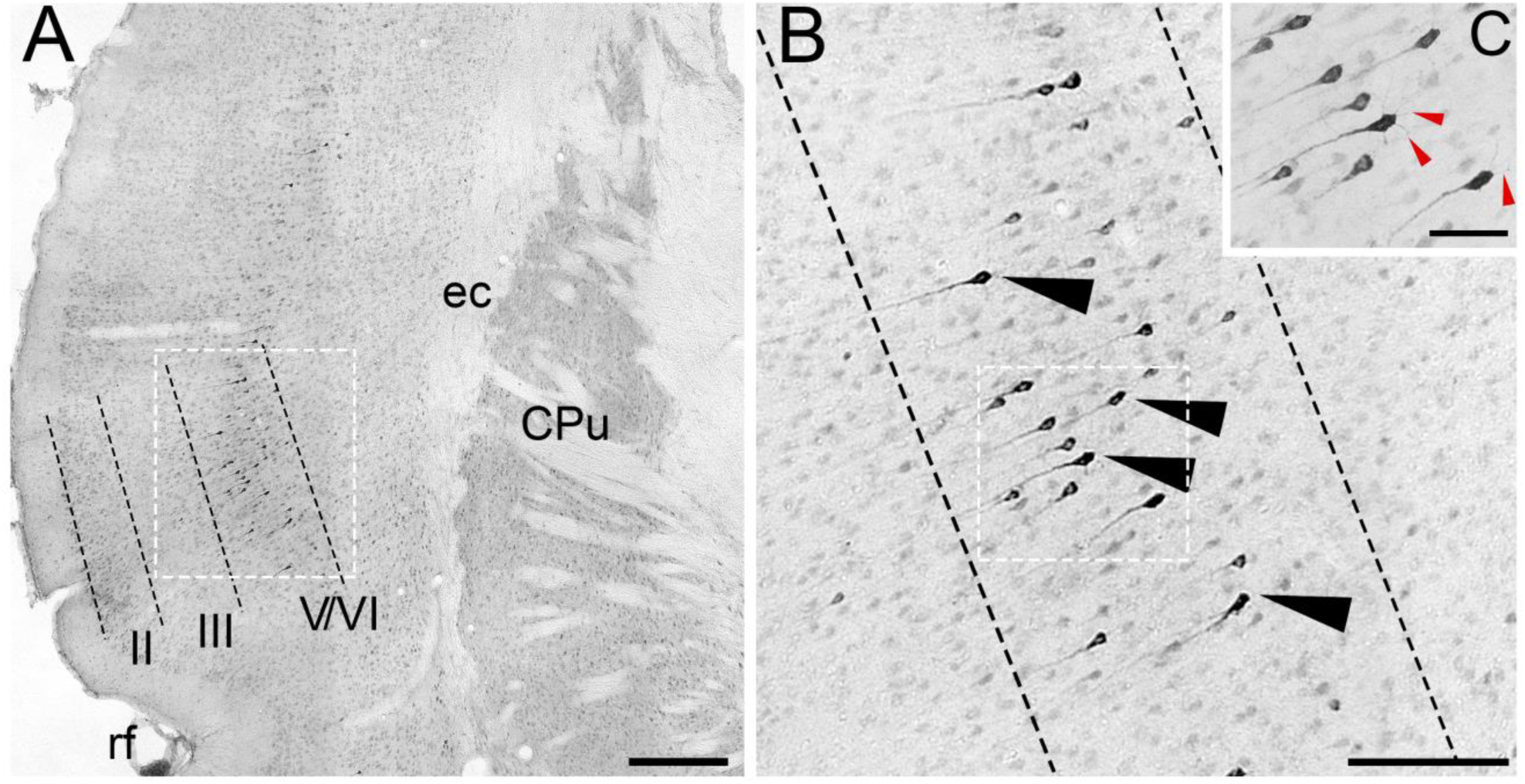
Representative CT-B-labeled pyramidal neurons in layer V/VI of the insular cortex (**A**). **B**&**C:** Photomicrographs show labeled neurons at progressively higher magnifications. **A**: the photomicrograph at the lowest magnification shows the specificity of retrograde labeling in the context of the neighboring cells and structures. **B**: at highest magnification, the morphology of labeled neurons (black arrowheads) is more apparent. **C**: at a slightly higher magnification than that in B, the dentrites (red arrowheads) of the pyramidal neurons are visible. *Abbreviations*: II, layer 2 of cortex ; III, layer 3 of cortex; V, layer 5 of cortex; CPu, caudate putamen (striatum); ec, external capsule; rf, rhinal fissure. *Scale bars*: A = 200 µm; B = 100 µm; C = 50 µm.

### 3.3. Specific distribution of retrogradely CT-B-labeled neurons in cortical and sub-cortical brain regions following injections in the midbrain periaqueductal gray (PAG)

Figure 4 shows representative images of retrogradely labeled neurons in the cortical and sub-cortical brain regions following PAG injections. Figure 5 illustrates the rostro-caudal gradients of all CT-B-labeled neurons in relation to bregma.

**Figure 4.**
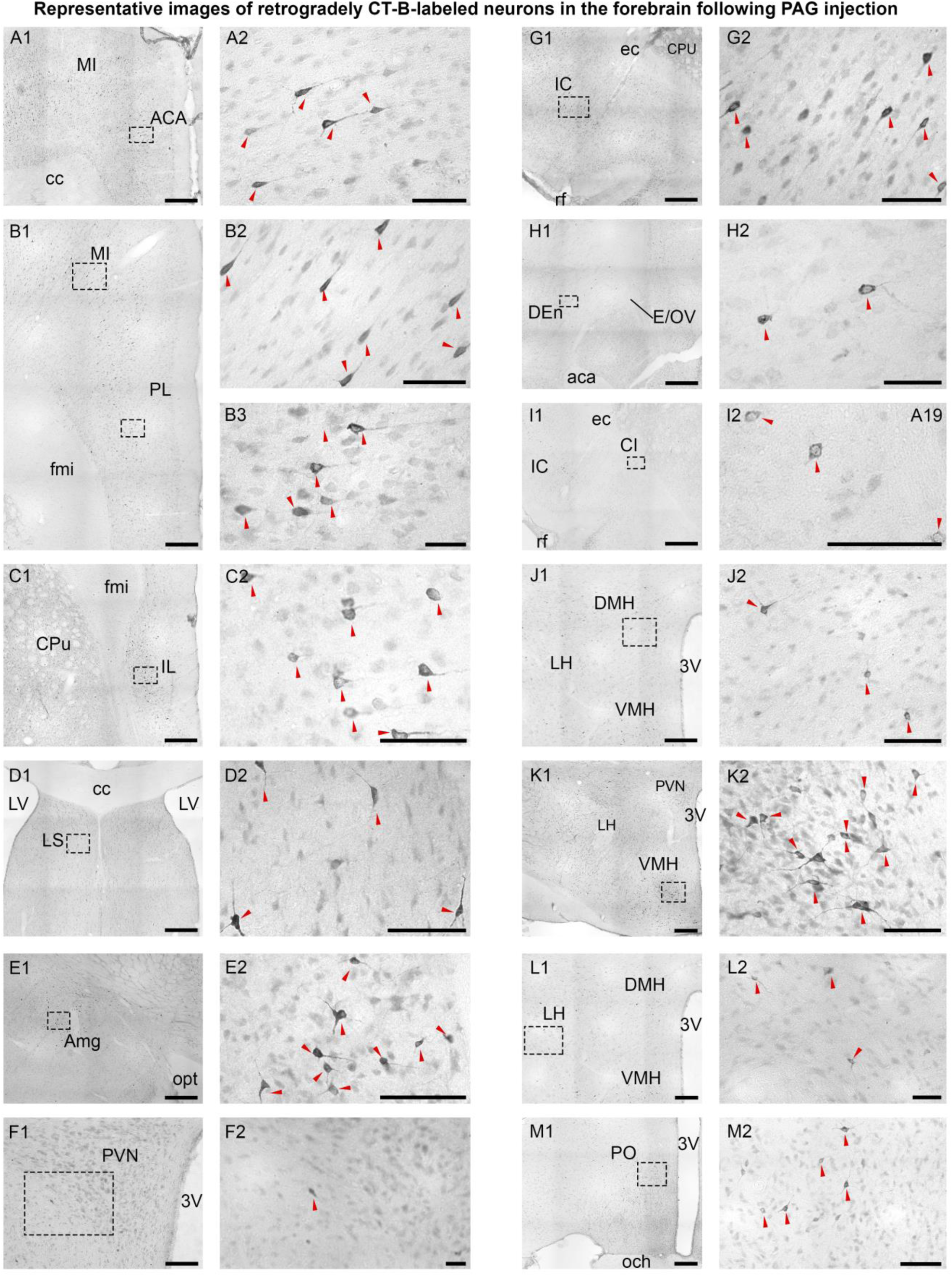
CT-B microinjection in the PAG labeled neurons in various cortical and forebrain regions. In this and figures 6, 8, & 11, we present the paired photomicrographs of the labeled neurons. **1**) The collage on the left is at low magnification to show the specificity of the retrolabeling in the context of the neighboring structures. Dashed-line boxes in 1 outline the area of collage in **2. 2**) The collage on the right is at high magnification to show the morphology of the labeled cells. Red arrowheads point to representative labeled neurons. The following areas had CT-B-labeled neurons: **A1-2**, the cingulate cortex (ACA); **B1-3**, motor and prelimbic cortices (MI, PL); **C1-2**, infralimibic (IL); **D1-2**, lateral septum (LS); **E1-2**, amygdala (Amg); **F1-2**, paraventricular hypothalamus (PVN); **G1-2**, insular cortex (IC); **H1-2**, endopiriform nucleus (DEn); **I1-2**, claustrum (CI); **J1-2**, dorsomedial (DMH); **K1-2**, ventromedial (VMH); **L1-2**, lateral hypothalmus (LH); and **M1-2**, preoptic area (PO). *Abbreviations*: 3V = third ventricle; aca = anterior commissure; cc = corpus callosum; CPu = caudate putamen (striatum); E/OV = ependymal and sublayer EV; ec = external capsule; fmi = forceps minor of corpus callosum; ic = internal capsule; LV = lateral ventricle; mlf = medial longitudinal fasciculus; och = optic chiasm; opt, optic tract; rf = rhinal fissure. *Scale bars* on this and Figures 6, 8, & 11: 200 µm (low-megnification images); 50 µm (high-magnification images).

**Figure 5.**
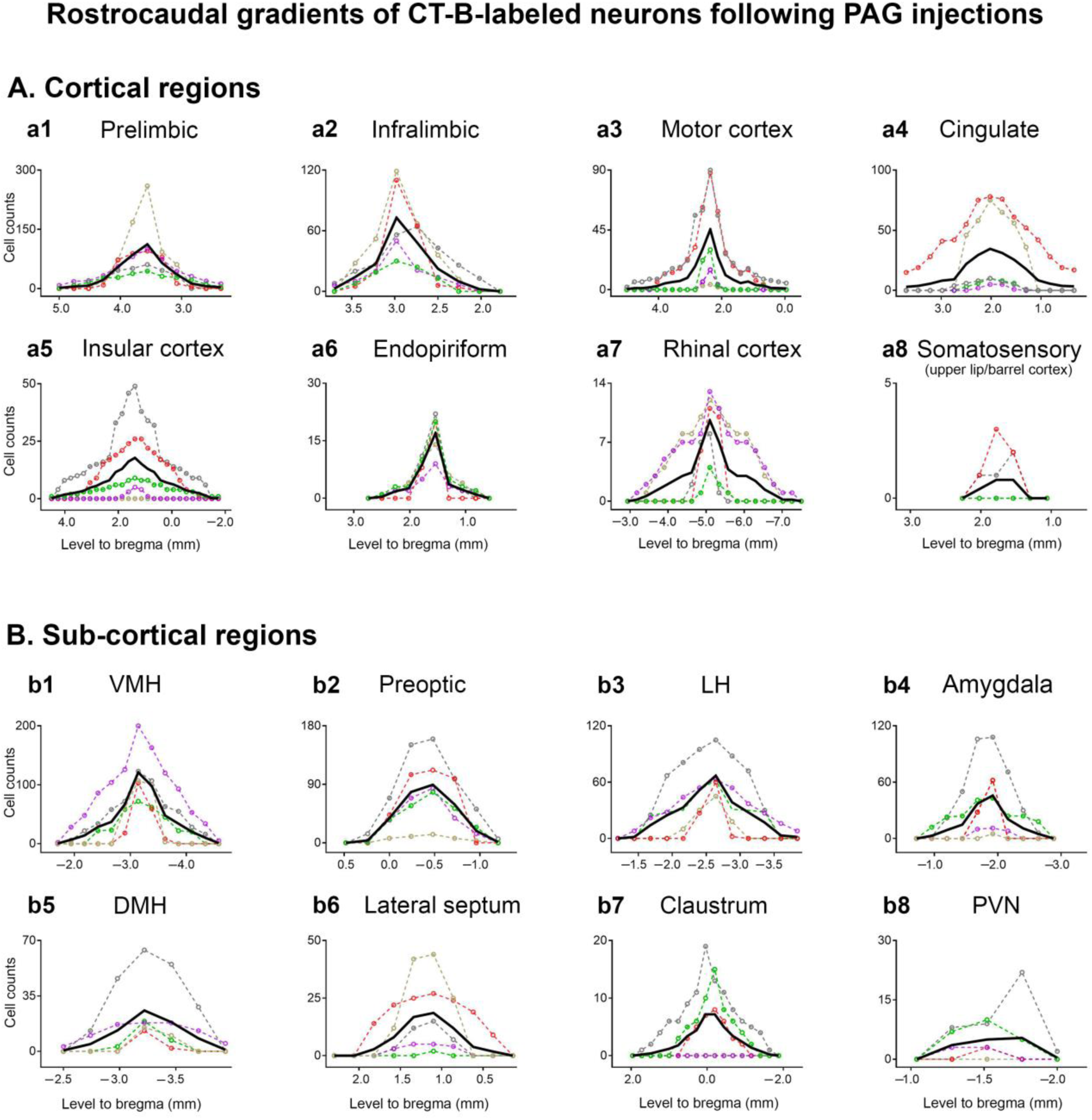
The rostrocaudal distribution of the numbers of retrogradely labeled neurons relative to bregma (mm) in various cortical (**A**) and sub-cortical (**B**) brain regions following CT-B microinjections in the periaqueductal gray. The location of the specific injection sites are represented by color-codes, which are the same as Figure 1. Black lines represent the mean number of CT-B-labeled neurons detected throughout the rostrocaudal levels for each cortical and sub-cortical brain region. *Abbreviations*: DMH = dorsomedial hypothalamus; LH = lateral hypothalamus; PAG = periaqueductal gray; PVN = paraventricular hypothalamus; VMH = ventromedial hypothalamus.

We observed the largest number of CT-B-labeled neurons in the prefrontal cortex (PFC), including the dorsal (Fig. 5a1; prelimbic, 412 ± 51 neurons/case [n/c]) and ventral PFC (Fig. 5a2 infralimbic, 198 ± 40 n/c). In prelimbic cortex, CT-B-labeled neurons occupied the rostral and intermediate levels (bregma: from +4.0 to +1.5 mm). In infralimbic cortex, CT-B-labeled neurons were predominantly located at rostral levels (bregma: from +4.0 to +2.5 mm). We also observed a large number of CT-B-labeled neurons in motor cortex (Fig. 5a3 ; 192 ± 99 n/c; bregma: from +4.0 to 0 mm). Substantial numbers of CT-B-labeled neurons in cingulate cortex were observed after injections in the rostral PAG (case #1, 336 neurons; case #2, 648 neurons), whereas injections in the caudal region of the midbrain PAG revealed smaller numbers of CT-B labeled-neurons in the cingulate cortex (case #3, 41 neurons; case #4, 33 neurons; case #5, 0 neurons) (Fig. 5). In addition, we also observed a considerable number of CT-B-labeled neurons in the insular cortex (Fig. 5a5; 172 ± 88 n/c, bregma: from +4.5 to +1.5 mm), particularly after injections in the rostral vlPAG (case #2, 272 neurons; case #3, 469 neurons). Caudal injections in the vlPAG (cases #4-5) revealed fewer numbers of CT-B-labeled neurons in the insular cortex (case #4, 14 neurons; case #5, 106 neurons), whereas no CT-B-labeled neurons were observed after injections in the rostral dlPAG (case #1). A small number of CT-B-labeled neurons were found in the rhinal (50 ± 20 n/c; bregma: from -3.18 to -7.3 mm) and endopiriform cortices (cases #1-5, 37 ± 4 n/c; Fig. 5a6;). CT-B-labeled neurons in the endopiriform were restricted to the rostral level of this nuclei (bregma: from -0.7 to +2.4 mm). Finally, smaller numbers of monosynaptic projections to the PAG were observed from the somatosensory cortex (2 ± 1 n/c; bregma: from +1.8 to +1.3 mm).

Substantial numbers of monosynaptic projections to the PAG were also observed from sub-cortical regions, such as the claustrum, amygdala, lateral septum and hypothalamus (Fig. 4 and 5). Injections in the PAG revealed high numbers of retrogradely labeled neurons in the amygdala (146 ± 72 n/c; bregma: from -1.0 to -2.6 mm), whereas a smaller number of labeled neurons were observed after injections in the dlPAG (case #1, 5 neurons). Injections in the rostral PAG revealed a large number of CT-B-labeled neurons in the lateral septum (case #1, 123 neurons; case #2, 140 neurons; bregma: from +0.6 to +1.8 mm), whereas injections in caudal PAG (case #4, 17 neurons; case #5, 2 neurons) yielded smaller numbers of descending projection neurons from the lateral septum. Hypothalamic nuclei showed high numbers of CT-B-labeled neurons in all cases (Fig. 5.b1, b2, b3, b5, b8). Overall, the highest numbers of retrogradely CT-B-labeled neurons were observed in the ventromedial hypothalamus (VMH, 486 ± 159 n/c; bregma: from -1.7 to -4.6 mm), followed by the preoptic area (PO, 288 ± 83 n/c; bregma: from -1.2 to +0.5 mm), lateral hypothalamus (LH, 280 ± 88 n/c; bregma: from -1.2 to -3.84 mm), dorsomedial hypothalamus (DMH, 72 ± 36 n/c; bregma: from -2.5 to -3.9 mm) and paraventricular hypothalamic nucleus (PVN, 14 ± 8 n/c; bregma: from -1.0 to -2.0 mm). Finally, a small number of CT-B-labeled neurons were observed in the claustrum after injections in the rostral or caudal PAG (40 ±2 0 n/c; bregma: from +2.2 to +1.6 mm). Overall, the distribution of forebrain projections to the midbrain PAG is consistent with the literature (Dampney et al., 2013; see discussion).

### 3.4. Specific distribution of retrogradely CT-B-labeled neurons in cortical and sub-cortical brain regions following injections in the pontine Kölliker-Fuse nucleus (KFn)

Figure 6 shows representative images of retrogradely CT-B-labeled neurons in the cortical and sub-cortical regions that project to the KFn. Figure 7 depicts the rostro-caudal gradients of CT-B-labeled neurons.

**Figure 6.**
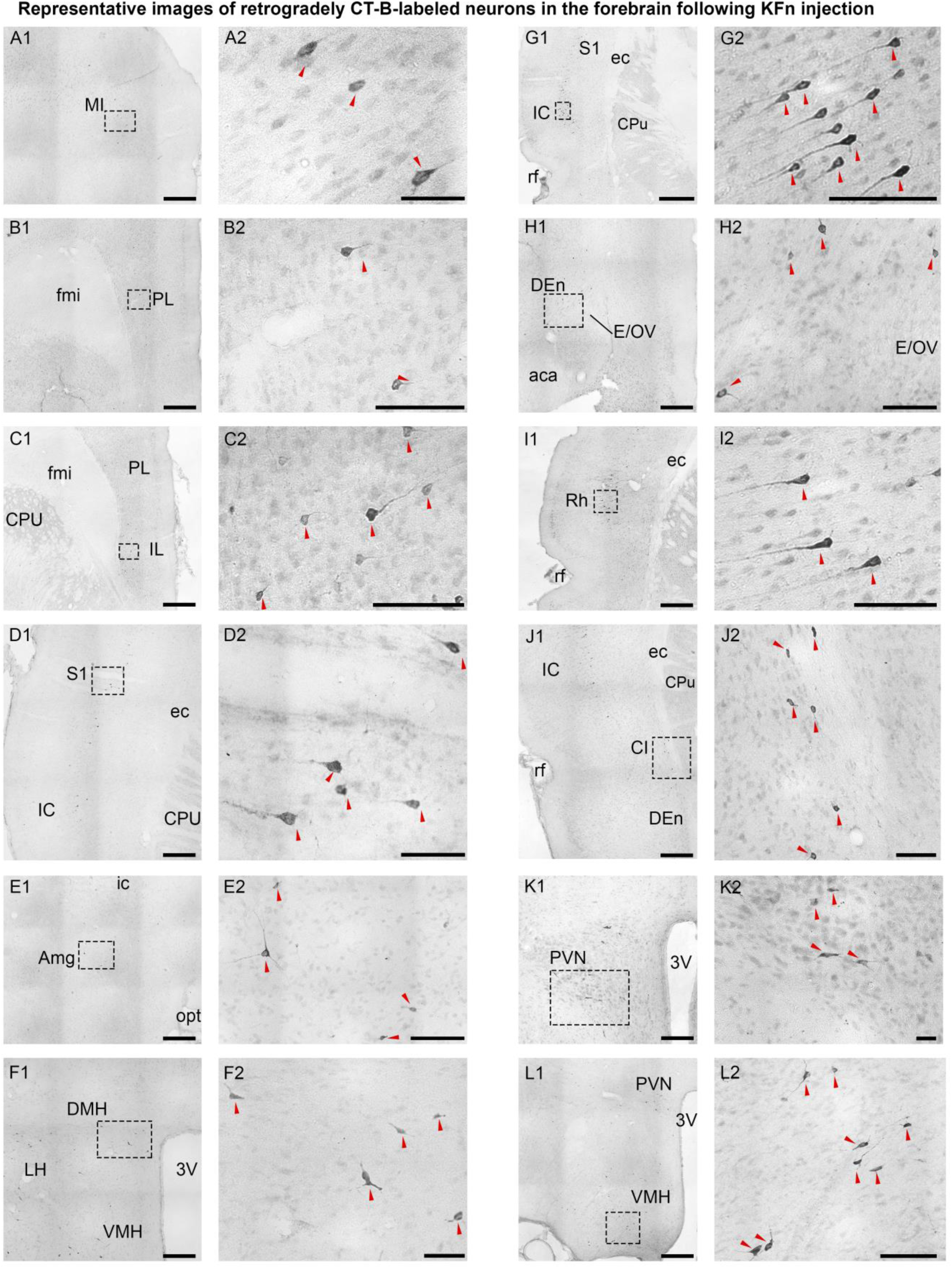
CT-B microinjection in the Kölliker-Fuse nucleus labeled neurons in various cortical and forebrain regions (paired photomicrographs: **1** is low and **2** is high magnification). The following areas had CT-B-labeled neurons: **A1-2**, motor cortex (MI); **B1-2**, pelimibc cortex (PL); **C1-2**, infralimibic (IL); **D1-2**, somtasensory cortex (S1); **E1-2**, amygdala (Amg); **F1-2**, dorsomedial hypothalamus (DMH); **G1-2**, insular cortex (IC); **H1-2**, endopiriform nucleus (DEn); **I1-2**, rhinal cortex (Rh); **J1-2**, claustrum (CI); **K1-2**, paraventricular hypothalamus (PVN); **L1-2**, ventromedial (VMH). *Abbreviations*: 3V = third ventricle; aca = anterior commissure; CPu = caudate putamen (striatum); E/OV = ependymal and sublayer EV; ec = external capsule; fmi = forceps minor of corpus callosum; ic = internal capsule; mlf = medial longitudinal fasciculus; och = optic chiasm; opt = optic tract; rf = rhinal fissure. *Scale bars*: 200 µm (low-magnification images); 50 µm (high-magnification images).

**Figure 7.**
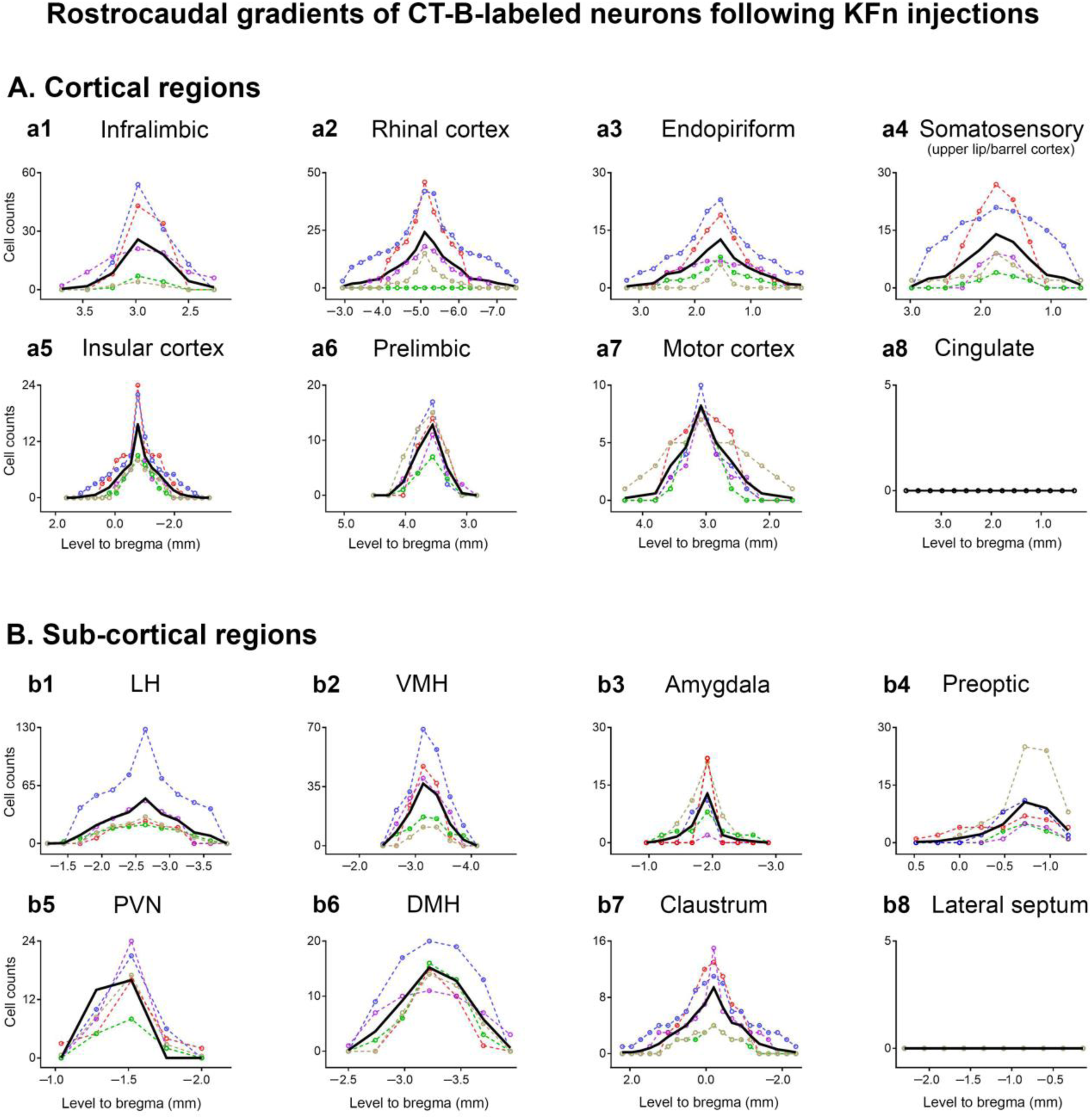
The rostrocaudal distribution of retrogradely labeled neurons relative to bregma (mm) in various cortical (**A**) and sub-cortical (**B**) brain regions following CT-B microinjections in the Kölliker-Fuse nucleus. The location of the specific injection sites are represented by color-codes, which are the same as figure 1. Black lines represent the mean number of CT-B-labeled neurons detected throughout the rostrocaudal levels for each cortical and sub-cortical brain region. *Abbreviations*: DMH = dorsomedial hypothalamus; KFn = Kölliker-Fuse; LH = lateral hypothalamus; PVN = paraventricular hypothalamus; VMH = ventromedial hypothalamus.

Quantitative analysis of monosynaptic projections to the KFn revealed considerable numbers of CT-B-labeled neurons in infralimbic (60 ± 21 n/c; bregma: from +3.7 to +2.3 mm), rhinal (146 ± 63 n/c; bregma: from -2.3 to -7.5 mm), endopiriform (60 ± 22 n/c; bregma: from +3 to +0.1 mm), somatosensory (63 ± 25 n/c; bregma: from +2.5 to +0.1 mm), insular (64 ± 15 n/c; bregma: from +1.4 to -2.5 mm), prelimbic (29 ± 5 n/c; bregma: from +4.0 to +3.7 mm), and motor cortices (29 ± 4 n/c; bregma: from +4.5 to +1.4 mm). In contrast to PAG injections, we did not detect CT-B-labeled neurons in the cingulate cortex following injections in the KFn.

Monosynaptic projections to the KFn were also found from sub-cortical areas such as the claustrum, amygdala and various hypothalamic nuclei. In contrast to PAG injections, no neurons were found in the lateral septum after CT-B injection in the KFn. The highest numbers of retrogradely labeled neurons were found in the LH (225 ± 88 n/c; bregma: from -1.44 to -3.6 mm), followed by the VMH (112 ± 33 n/c; bregma: from -2.7 to -3.6 mm), DMH (47 ± 8 n/c; bregma: from -2.5 to -3.9 mm), PO (34 ± 12 n/c; bregma: from +0.5 to -1.2 mm), and PVN (29 ± 4 n/c; bregma: from -1.3 to -2 mm). We also observed CT-B-labeled neurons in the claustrum (Fig. 7b7; 56 ± 1 n/c; bregma: from +2.2 to -2.4 mm) and amygdala (23 ± 7 n/c; bregma: from -1.2 to -2.6 mm).

### 3.5. Specific distribution of retrogradely CT-B-labeled neurons in cortical and sub-cortical brain regions following injections in the medullary pre-Bötzinger complex (pre-BötC) and Bötzinger complex (BötC)

Figure 8 shows representative images of retrogradely CT-B-labeled neurons in the cortical and sub-cortical regions following CT-B injections in the pre-BötC. Figures 9 (pre-BötC injections) and 10 (BötC injections) illustrate the rostro-caudal gradients of CT-B-labeled neurons in relation to bregma.

**Figure 8.**
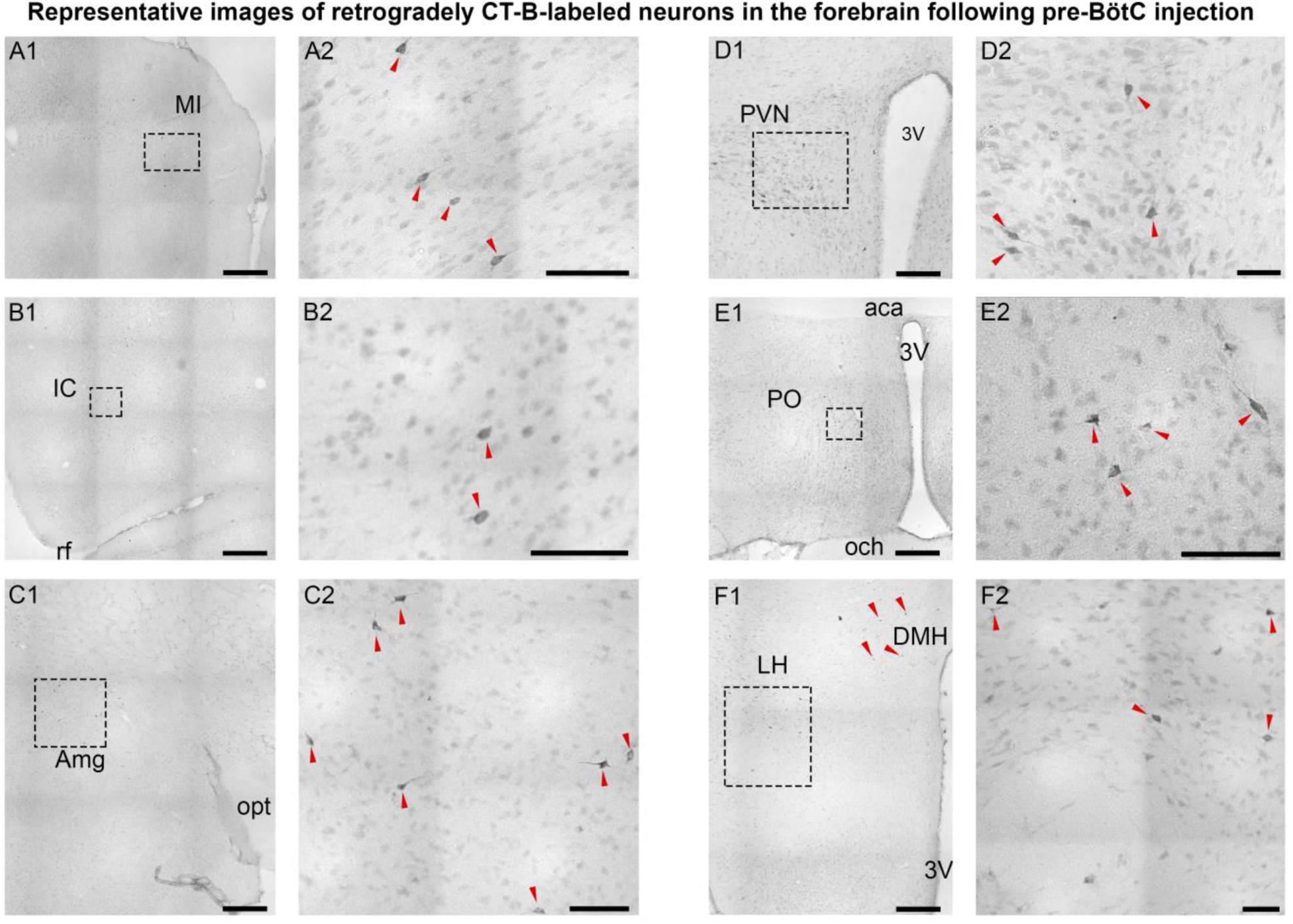
CT-B microinjection in the pre-Bötzinger complex labeled neurons in various cortical and forebrain regions (paired photomicrographs: **1** is low and **2** is high magnification). The following areas had CT-B-labeled neurons: **A1-2**, motor cortex (MI); **B1-2**, insular (IC); **C1-2**, amygdala (Amg); **D1-2**, paraventricular hypothalamus (PVN); **E1-2**, preoptic area (PO); and **F1-2**, lateral hypothalamus (LH). *Abbreviations*: 3V = third ventricle; och = optic chiasm; pre-BötC = pre-Bötzinger complex; rf = rhinal fissure. *Scale bars*: 200 µm (low-magnification images); 50 µm (high-magnification images).

**Figure 9.**
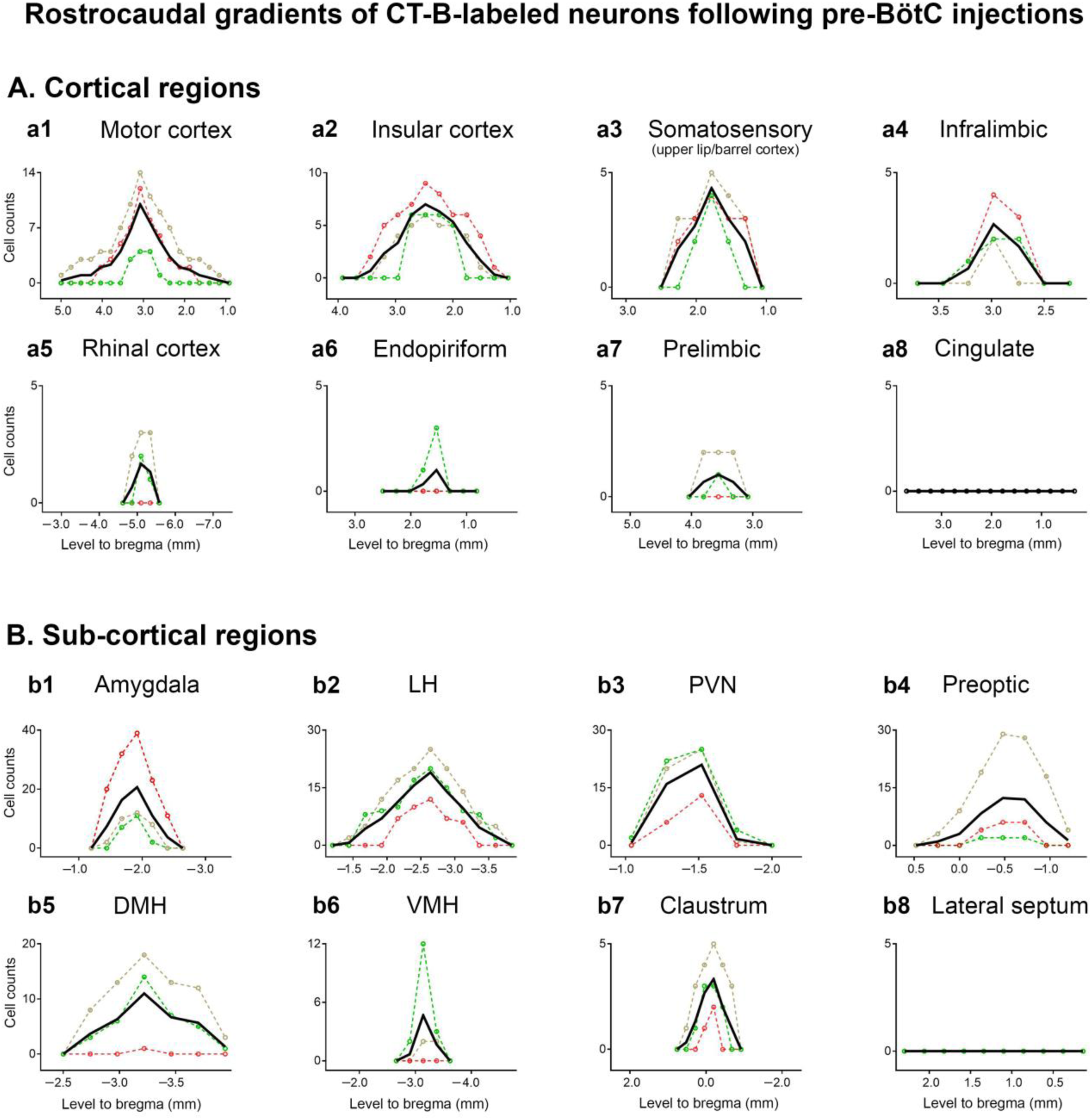
The rostrocaudal distribution of retrogradely labeled neurons relative to bregma (mm) in various cortical (**A**) and sub-cortical (**B**) brain regions following CT-B microinjections in the pre-Bötzinger complex. The location of the specific injection sites are represented by color-codes, which are the same as Figure 1. Black lines represent the mean numbers of CT-B-labeled neurons detected throughout the rostrocaudal levels for each cortical and sub-cortical brain region. *Abbreviations:* DMH = dorsomedial hypothalamus; LH = lateral hypothalamus; pre-BötC = pre-Bötzinger complex; PVN = paraventricular hypothalamus; VMH = ventromedial hypothalamus.

The highest number of descending projection neurons to the pre-BötC was observed in the motor cortex (50 ± 21 n/c; bregma: from +5 to +1.2 mm), followed by the insular (36 ± 9 n/c; bregma: from +3.4 to +1.3 mm), somatosensory (14 ± 3 n/c; bregma: from +2 to +1.1 mm), infralimbic (5 ± 2 n/c; bregma: from -3.2 to -2.7 mm), rhinal (4 ± 2 n/c; bregma from - 5.3 to -4.9 mm), prelimbic (2 ± 2 n/c; bregma: from +3.8 to +3.3 mm), and endopiriform cortices (1 ± 1 n/c; bregma: from +1.7 to +1.4 mm). Similar to KFn CT-B injections, we did not detect labeled neurons in the cingulate cortex after injections in the medullary pre-BötC.

In sub-cortical regions (Fig. 9), we observed monosynaptic projections to the pre-BötC from amygdala (59 ± 33 n/c; bregma: from -1.4 to -2.4 mm), claustrum (11 ± 5 n/c; bregma: from +0.5 to -0.7 mm) and hypothalamic nuclei, including the LH (89 ± 25 n/c; bregma: from -1.4 to -3.6 mm), PO (45 ± 34 n/c; bregma: from +0.3 to -1.4 mm), PVN (39 ± 10 n/c; bregma: from -2 to -1 mm), DMH (35 ± 19 n/c; bregma: from -2.7 to -3.9 mm), and VMH (7 ± 5 n/c; bregma: from -2.9 to -3.4 mm).

By contrast, CT-B injections in the BötC of the ventral respiratory column revealed substantially fewer numbers of descending projection neurons in cortical brain regions (p=0.0003). We observed CT-B-labeled neurons projecting to BötC in the insular (31 ± 28 n/c; bregma: from +3.7 to +1.3 mm), endopiriform (3 ± 2 n/c; bregma: from +1.9 to +1.2 mm), motor (1 ± 1 n/c; bregma: +3.3 mm), infralimbic (1 ± 1 n/c; bregma: +3.0 mm), somatosensory (1 ± 1 n/c; bregma: +1.6 mm), rhinal (1 ± 1 n/c; bregma: from -5.1 to -5.4 mm) and prelimbic cortices (1 ± 1 n/c; bregma: +3.6 mm). Similarly, only small numbers of CT-B-labeled neurons were detected in sub-cortical regions following BötC injections: amygdala (21 ± 21 n/c; bregma: from -1.7 to -2.2 mm), LH (3 ± 2 n/c; bregma: from -2.4 to - 2.6 mm), PO (1 ± 1 n/c; bregma: -0.5 mm), PVN (1 ± 1 n/c; bregma: -1.5 mm), and claustrum (1 ± 1 n/c; bregma: -0.2 mm). Finally, CT-B-labeled neurons in lateral septum, DMH and VMH were never observed after injections in the medullary BötC.

### 3.6. Specific distribution of retrogradely CT-B-labeled neurons in cortical and sub-cortical brain regions following injections in the caudal raphé in the medulla oblongata

Figure 11 shows representative images of retrogradely CT-B-labeled neurons in cortical and sub-cortical regions following CT-B injections in the caudal raphé nuclei of the medulla oblongata. Figure 12 illustrates the quantitative analysis of CT-B-labeled neurons along their rostrocaudal gradients in relation to bregma. In accordance with published literature (Hermann et al., 1997), CT-B injections in the caudal raphé showed variable numbers of labeled neurons in the forebrain, which is dependent whether the injections were centred in the RMg, ROb, or RPa. For example, case #17 (injection in the caudal RPa) revealed the highest number of CT-B-labeled neurons in the cortex (Fig. 12).

**Figure 10.**
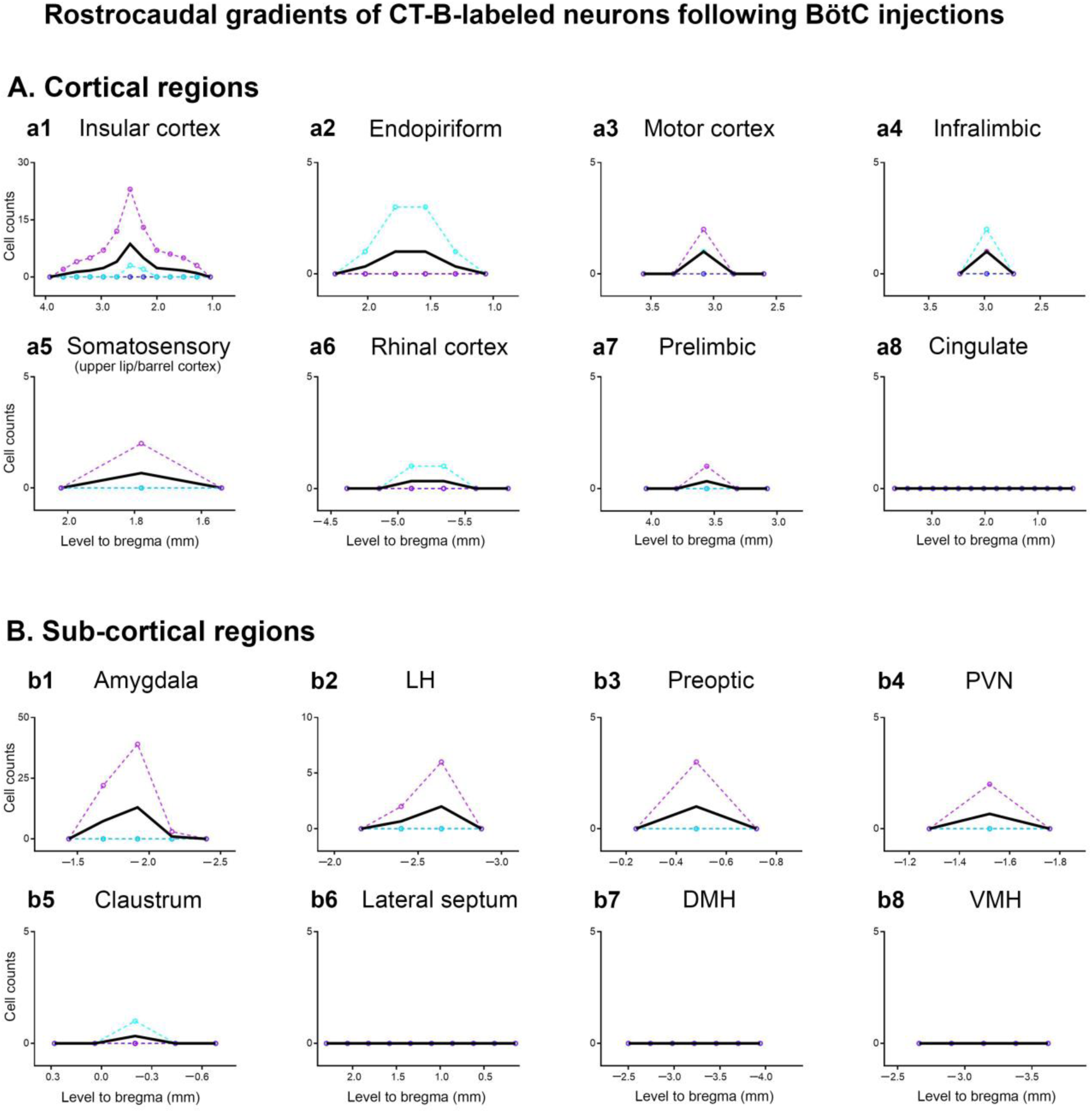
The rostrocaudal distribution of the numbers of retrogradely labeled neurons relative to bregma (mm) in various cortical (**A**) and sub-cortical (**B**) brain regions following CT-B microinjections in the Bötzinger complex. The location of the specific injection sites are represented by color-codes, which are the same as Figure 1. Black lines represent the mean number of CT-B-labeled neurons detected throughout the rostrocaudal levels for each cortical and sub-cortical brain region. *Abbreviations*: BötC = Bötzinger complex; DMH = dorsomedial hypothalamus; LH = lateral hypothalamus; PVN = paraventricular hypothalamus; VMH = ventromedial hypothalamus.

**Figure 11.**
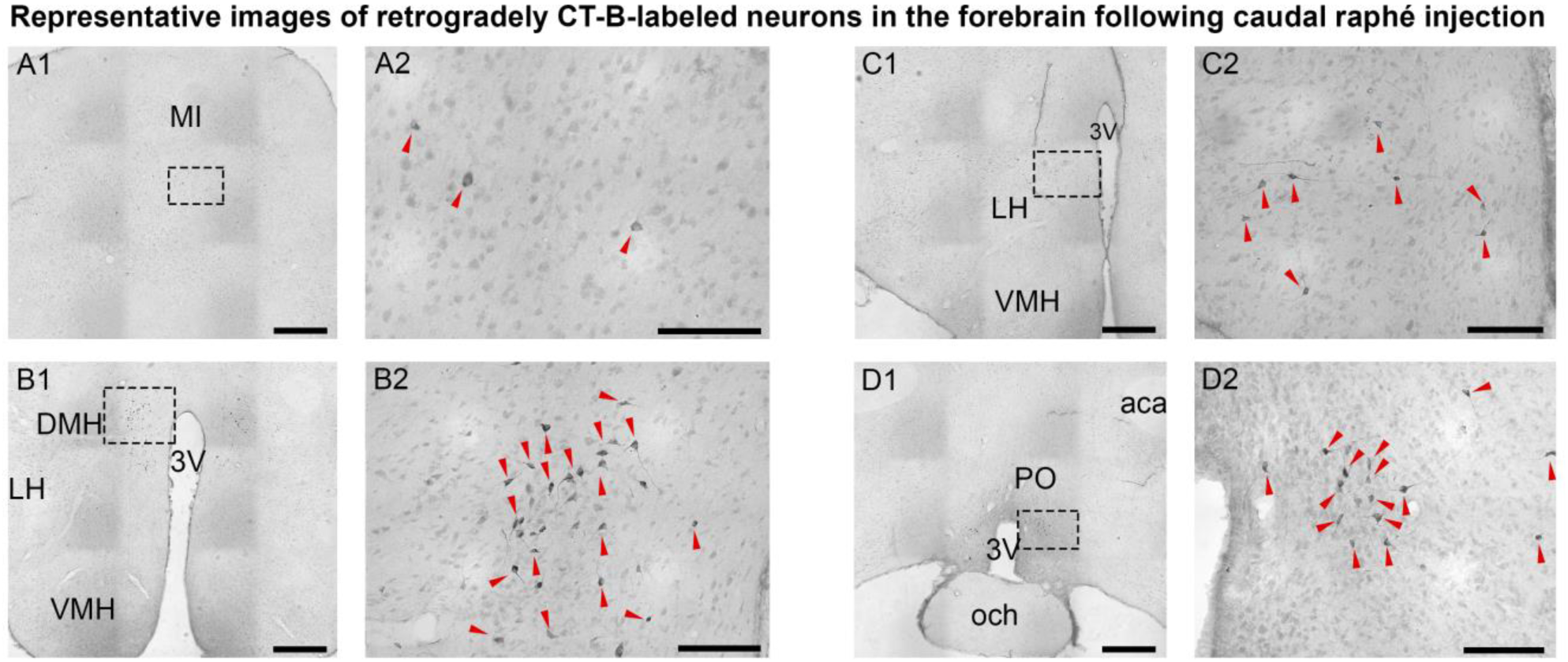
CT-B microinjection in the caudal raphé nuclei labeled neurons in various cortical and forebrain regions (paired photomicrographs: **1** is low and **2** is high magnification). The following areas had CT-B-labeled neurons: **A1-2**, motor cortex (MI); **B1-2**, dorsomedial hypothalamus (DMV); **C1-2**, lateral hypothalamus (LH); and **D1-2**, preoptic area (PO). *Abbreviations:* 3V = third ventricle; aca = anterior commissure; DMH, dorsomedial hypothalamus; LH, lateral hypothalamus; MI, motor cortex; och, optic chiasm; PO, preoptic nucleus; VMH, ventromedial hypothalamus. *Scale bars*: 200 µm (low-magnification images); 50 µm (high-magnification images).

**Figure 12.**
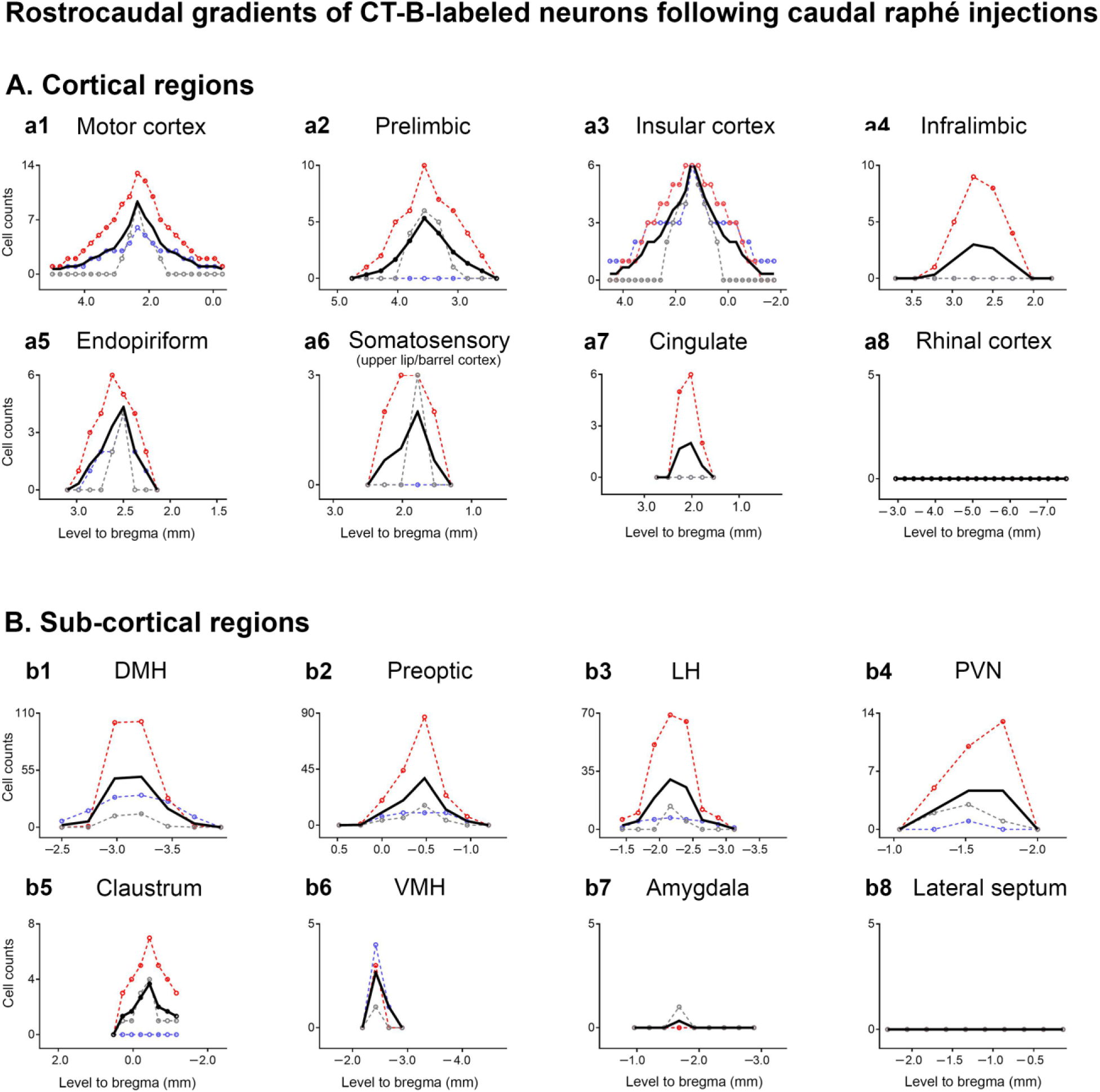
The rostrocaudal distribution of retrogradely labeled neurons relative to bregma (mm) in various cortical (**A**) and sub-cortical (**B**) brain regions following CT-B microinjections into the caudal raphé nuclei. The location of the specific injection sites are represented by color-codes, which are the same as Figure 1. Black lines represent the mean number of CT-B-labeled neurons detected throughout the rostrocaudal levels for each cortical and sub-cortical brain region. *Abbreviations*: DMH = dorsomedial hypothalamus; LH = lateral hypothalamus; PVN = paraventricular hypothalamus; VMH = ventromedial hypothalamus.

CT-B injections in the caudal raphé revealed a modest number of monosynaptic projections from cortex. The highest numbers of descending projection neurons were observed in the motor (66 ± 26 n/c; bregma: from +5 to +0.3 mm), followed by the insular (59 ± 13 n/c; bregma: +4.5 to +1.7 mm), prelimbic (20 ± 12 n/c; bregma: from +5 to +1.9 mm), endopiriform (14 ± 6 n/c; bregma: from +2.6 to +1.2 mm), infralimbic (9 ± 9 n/c; bregma: from +3.2 to +2.2 mm), somatosensory (4 ± 3 n/c; bregma: from +3 to +1.6 mm), and cingulate cortices (4 ± 4 n/c; bregma: from +2.3 to +1.8 mm). No CT-B-labeled neurons were detected in the rhinal cortex followed by injections in any caudal raphé nucleus.

In sub-cortical regions, descending connectivity outside the hypothalamus was absent or weak (e.g. claustrum, amygdala and lateral septum). However, high to moderate numbers of projection neurons were observed in various hypothalamic nuclei. The highest numbers of CT-B-labeled neurons were in the DMH (125 ± 60 n/c; bregma: from -2.5 to -3.7 mm), followed by the LH (91 ± 64 n/c; bregma: from -1.4 to -3.1 mm), and PO (84 ± 49 n/c; bregma from +0.2 to -1 mm). The number of CT-B-labeled neurons in the PVN (12 ± 8 n/c; bregma: from -1.8 to -1.3 mm) and VMH (3 ± 1 n/c; bregma: from -2.2 to -2.7 mm) were smaller compared to the aforementioned hypothalamic nuclei.

### 3.7 CT-B-labeled neurons associated with near-miss injections

Analysis of near-miss injection sites (*n*=9) presented far fewer labeled neurons compared to injections that were centered in the target areas. For instance, injections placed ventrolateral to the PAG presented far fewer labeled neurons in the forebrain. Only small numbers of CT-B-labeled neurons were observed, predominantly in the insular, rhinal and motor cortices, but not in hypothalamic areas (data not shown). In contrast, near-miss injections ventrocaudally to the KFn revealed modest numbers of CT-B-labeled neurons in the cortex, but we still identified robust numbers of projection neurons in the hypothalamus (e.g. lateral hypothalamus) and midbrain PAG (data not shown). The later illustrates that the caudal extension of the KFn remains connected to key nuclei of the autonomic nervous system in the forebrain, but lacks major cortical inputs. Finally, near-miss injections placed in the reticular formation outside the pre-BötC, BötC and caudal raphé target areas resulted in almost complete absence of forebrain projection neurons, although labeled cells were still observed in the KFn and PAG (data not shown). Thus, the near miss injections underline the specificity of the descending projection patterns in the investigated respiratory nuclei (as well as the specificity of CT-B antibody used in this study).

## 4. Discussion

We identified wide-spread monosynapic descending projections from the forebrain to all respiratory control areas investigated suggesting that forebrain-evoked modulations of respiration do not simply bypass brainstem circuits, but instead, are likely co-ordinated with spontaneous respiratory network activity. Descending projection neurons that target designated respiratory control areas in the midbrain and brainstem were restricted to a variety of cortical areas, the claustrum, lateral septum, various hypothalamic nuclei and the amygdala. Examination of the distribtution of retrogradely labeled neurons across the entire axis of the forebrain failed to detect any descending projection neurons in some major neural systems of the forebrain, such as the thalamus, hippocampus, olfactory bulb or basal ganglia.

The topographical organization of detected mono-synatpic descending forebrain projections is summarized in a network connectivity graph (Figure 13). The graph shows the relative proportion of projections from a given cortical or sub-cortical area to each respiratory control nuclei investigated. The general distribution of descending projection neurons located in sub-cortical areas, such as the hypothalamus or amygdala, reveals a broad connectivity pattern with the downstream targets. These sub-cortical projections are discussed in the context of their putative role in homeostastis, state-dependent and emotional modulation of breathing (section 4.1). In contrast, the cortical descending connectivity measured in the present study implies that retrogradely labeled neurons predominantly target the midbrain PAG and pontine KFn. The only exception is the insular cortex which also provides an equal proportion of descending inputs to all investigated respiratory control areas. The long-range forebrain-brainstem projection neurons detected in this study were exclusively located in the deepest layers of cortex. The role of these cortex-brainstem projections is discussed in the context of volitional control of breathing (Section 4.2).

**Figure 13.**
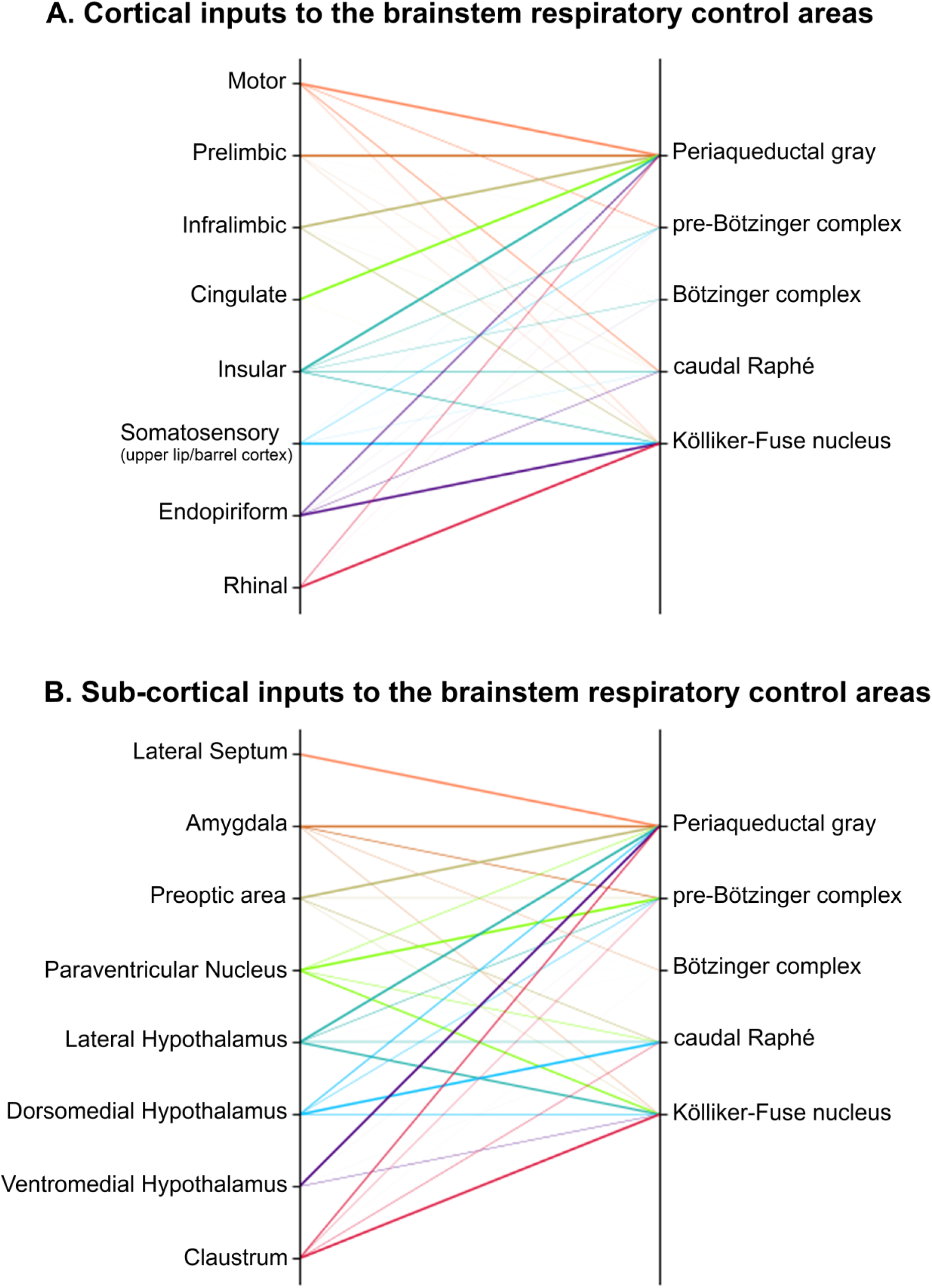
Summary of the relative strength of descending projections arising from cortical **(A)** and sub-cortical **(B)** areas in a connectivity map. The weight of connecting lines are proportional to the normalized maximal number of CT-B-labeled neurons found in the specific source of descending projections.

Finally, amongst all downstream targets investigated, the BötC shows the least amount of forebrain connectivity, despite its designated key function in respiratory pattern formation (Burke et al., 2010; Smith et al., 2013; Marchenko et al., 2016).

### 4.1. Homeostasis, state-dependent or emotional modulation of respiration

The dense descending connectivity of visceral sensory areas of the insular cortex with respiratory control areas confirm previous anatomical studies (Saper, 1982; Sato et al., 2013; Grady et al., 2020), and are in line with several functional studies that have associated the insular cortex with cardio-respiratory modulation (Rugiero et al., 1987; Yasui et al., 1991; Aleksandrov et al., 2000), which may be associated with adaptation of breathing to interoceptive states (Verdejo-Garcia et al., 2012). The present study also confirms well-documented descending projections of hypothalamic nuclei targeting the PAG, KFn, pre-BötC, BötC and caudal raphé (Peyron et al., 1998; Geerling et al., 2010). Indeed, a range of respiratory patterns, in the context of homeostasis, state-dependency (sleep-wake), and stress, can be evoked by hypothalamic subnuclei (for review, see Fukushi et al., 2019).

The insular cortex and hypothalamus are part of the widely-distributed limbic system that includes the amygdala, and the piriform, rhinal and prefrontal cortices. Because emotions have a profound influence on respiratory activity (Harper et al. 1984; Onimaru and Homma, 2007; Homma and Masaoka, 2008; Holstege, 2014; Subramanian and Holstege, 2014), it is not surprising that all aforementioned areas have direct descending projections to several respiratory control areas investigated in the present study (Fig.13). While it was previously postulated that the midbrain PAG, with its known function in respiratory modulation in relation to fear and defensive behavior (Carrive et al., 1988; Depaulis et al., 1989; Carrive, 1993; Carrive et al., 1997; Subramanian and Holstege, 2014), is the primary interface that links emotion with breathing, the present study illustrates that similar descending inputs also target nuclei of the primary respiratory rhythm and pattern generating network in the brainstem. Significant inputs from the amygdala, and the pre- and infra-limbic and rhinal cortices to the pontine KFn and medullary pre-BötC, with their specific function in controling respiratory phase transitions (Dutschmann and Herbert 2006) and rhythm generation (Smith et al., 1991; Feldman and Del Negro, 2006; Del Negro et al., 2018), suggests that the synaptic output of the widely-distrubted limbic system also connects to a similarly distributed respiratory network spanning from the midbrain to medulla oblongata. Considering recent evidence suggesting that action/motor encoding involves cell assemblies whose members are distributed in many areas beyond the cortex, including the PAG (Steinmetz et al., 2019), the present study supports the working hypothesis that the precise encoding of homeostatic, state-dependent or emotional breathing patterns may be partially outsourced to the respiratory network.

### 4.2 Volitional control of respiration

In contrast with previous suggestions that cortical motor commands for the respiratory system may be mediated via cortico-spinal pathways that connect the cortex with primary respiratory motor pools (Rikard-Bell et al., 1985; Gandevia and Rothwell, 1987a,b; Pouget et al., 2008), our study identifies distinct descending pathways that connect cortical output neurons with specific respiratory pre-motor nuclei (Fig. 13A). For instance, descending inputs from cortical areas of the frontal lobe, including the PFC (pre- and infra-limbic cortices) and the cingulate cortex, have the strongest connectivity with the lateral and ventrolateral columns of the PAG.

Amongst all cortical and sub-cortical areas with descending projections to respiratory control areas, the cingulate and lateral septum almost exclusively target the PAG (Fig. 13A). The neural networks composed of the cingulate and septal nuclei, including the prefrontal cortex (area of Brocca), harbor the primary synaptic network for the generation of speech in humans and vocalization in mammals (Jürgens, 2009; Hage and Nieder, 2016; Holstege and Subramanian, 2016). In line with the specific descending projection pattern that links the PAG to volitional motor commands for vocalization, stimulation of the midbrain PAG (particularly the lPAG and vlPAG) evokes vocalization in animals (Larson, 1985; Jürgen and Richter, 1986; Bandler and Carrive, 1988; Zhang et al., 1994). Thus, the present study supports the hypothesis that the midbrain PAG is the “final common pathway” for vocalization in mammals (Jürgens and Richter, 1986; Holstege, 1989; Jürgens 1994; Jürgens 2002; Düsterhöft et al., 2004; Subramanian and Holstege, 2009).

Studies in humans, however, have shown that the midbrain PAG is not activated during other voluntarily controlled orofacial behaviors such as coughing (Mazzone et al., 2011) and swallowing (Zald et al., 1999). Morover, the PAG neither possesses direct synaptic connection to cranial and spinal motor respiratory pools (Holstege, 1989; Zhang et al., 1995; Subramanian and Holstege, 2009; Dampney, 2013; Holstege, 2014), nor is it part of the primary rhythm and pattern generating circuit (Farmer et al., 2014). Thus, the midbrain PAG, as a mediator for modulation or reconfiguration of respiratory activity during orofacial behaviors, arousal or emotion, requires additional downstream synaptic interactions with the ponto-medullary brainstem respiratory network.

The PAG shares the highest level reciprocal connectivity with the KFn when compared to other medullary nuclei of the primary respiratory network (BötC, pre-BötC, or caudal raphé; data not shown, Trevizan-Baú et al., 2020, unpublished). Since both nuclei receive dense innervation from the claustrum, a general cortico-thalamic integration and output nucleus (Dillingham et al., 2017), it is possible that the PAG and the KFn may mediate various volitional motor commands in concert.

However, respiratory functions of the KFn in the mediation of the inspiratory-expiratory phase transition (Caille et al., 1981; Wang et al., 1993; Smith et al., 2007; Mörschel and Dutschmann, 2009) and gating of the respiratory cranial nerve activites (Dutschmann and Herbert, 2006 Bautista and Dutschmann, 2014), including in the branch of the facial nerve that innervates the nostrils and whisker pads (Dutschmann et al., 2020, unpublished), prime the KFn as a target for volitional motor commands for orofacial behavior. For instance, laryngeal adduction and the coordination of facial, vagal, and hypoglossal nerve activity are prerequisites for the mediation of swallowing (Dick et al., 1993; Bautista and Dutschmann, 2014) and sniffing (Semba et al., 1986; Perez Lobes, 2016). The present study now identifies significant descending input from the piriform and rhinal cortices to the KFn, suggesting that the KFn nucleus might receive major primary olfactory information (Chapuis et al., 2013; Leitner et al., 2016; Blazing and Franks, 2020). Additional substantial descending inputs to the KFn originate from the somatosensory barrel cortex, implying that the KFn also receives tactile sensory information from the whisker pads (Woolsey and Van der Loos, 1970; Petersen, 2019). In summary, because the KFn gates facial and upper-airway respiratory motor activity (Dutschmann et al., 2020, unpublished) and also receives cortical mono-synaptic tactile and olfactory sensory inputs, it is tempting to postulate that the KFn mediates volitional motor commands to coordinate breathing activity with sniffing and whisking.

### 4.3 Conclusions

In conclusion, the descending projection pattern of neurons of the widely-distributed limbic network (i.e. amygdala, rhinal cortex, endopiriform and the prefrontal cortex) connects to a similarly distributed network of respiratory control neurons from the midbrain to ponto-medullary brainstem. However, it is likely that specific volitional motor commands for vocalization are mediated via the midbrain PAG, while the coordination of whisking, sniffing, and swallowing with breathing activity is mediated by inputs that target the KFn. Amongst the medullary respiratory nuclei, the pre-BötC and caudal raphé receive limited descending input from the cortex (e.g. motor cortex), and therefore may have some additional function in the mediation or synaptic priming of volitional motor commands.

## Acknowledgements

We acknowledge that this work was conducted on the traditional land of the Wurundjeri people of the Kulin nation. We pay our respect to their elders past, present and emerging. This work was supported by research grants from the National Health and Medical Research Council of Australia (APP1165529 to MD & DS, and APP1078943 to SM); and Australian Research Council (Discovery project DP170104861); PT-B is funded by Melbourne Research Scholarship (University of Melbourne; 181858).

## Notes

**Conflict of interest** The authors have no conflicts of interest to declare.

### Competing Interest Statement

The authors have declared no competing interest.

## References

Aleksandrov, V. G., Aleksandrova, N. P., & Bagaev, V. A. (2000). Identification of a respiratory related area in the rat insular cortex. Canadian Journal of Physiology and Pharmacology, 78(7), 582–586. https://doi.org/10.1139/y00-031

Bandler, R., & Carrive, P. (1988). Integrated defence reaction elicited by excitatory amino acid microinjection in the midbrain periaqueductal grey region of the unrestrained cat. Brain Research, 439(1-2), 95–106. https://doi.org/10.1016/0006-8993(88)91465-5

Bandler R, Carrive P, & Depaulis A. (1991). Emerging principles of organization of the midbrain periaqueductal gray matter. In The midbrain periaqueductal gray matter (pp. 1–8). Springer, Boston, MA.

Bautista, T. G., & Dutschmann, M. (2014). Ponto-medullary nuclei involved in the generation of sequential pharyngeal swallowing and concomitant protective laryngeal adduction in situ. The Journal of Physiology, 592(12), 2605–2623. https://doi.org/10.1113/jphysiol.2014.272468

Besnard, S., Denise, P., Cappelin, B., Dutschmann, M., & Gestreau, C. (2009). Stimulation of the rat medullary raphe nuclei induces differential responses in respiratory muscle activity. Respiratory Physiology & Neurobiology, 165(2-3), 208–214. https://doi.org/10.1016/j.resp.2008.12.004

Blazing, R. M., & Franks, K. M. (2020). Odor coding in piriform cortex: mechanistic insights into distributed coding. Current Opinion in Neurobiology, 64, 96–102. https://doi.org/10.1016/j.conb.2020.03.001

Burke, P. G. R., Abbott, S. B. G., McMullan, S., Goodchild, A. K., & Pilowsky, P. M. (2010). Somatostatin selectively ablates post-inspiratory activity after injection into the Bötzinger complex. Neuroscience, 167(2), 528–539. https://doi.org/10.1016/j.neuroscience.2010.01.065

Butler, J. E. (2007). Drive to the human respiratory muscles. Respiratory Physiology & Neurobiology, 159(2), 115–126. https://doi.org/10.1016/j.resp.2007.06.006

Caille, D., Vibert, J. F., & Hugelin, A. (1981). Apneusis and apnea after parabrachial or Kölliker-Fuse N. lesion; influence of wakefulness. Respiration Physiology, 45(1), 79–95. https://doi.org/10.1016/0034-5687(81)90051-7

Carrive, P., Bandler, R., & Dampney, R. A. L. (1988). Anatomical evidence that hypertension associated with the defence reaction in the cat is mediated by a direct projection from a restricted portion of the midbrain periaqueductal grey to the subretrofacial nucleus of the medulla. Brain Research, 460(2), 339–345. https://doi.org/10.1016/0006-8993(88)90378-2

Carrive, P. (1993) The periaqueductal gray and defensive behavior: functional representation and neuronal organization. Behavioural Brain Research, 58, 27–47. https://doi.org/10.1016/0166-4328(93)90088-8

Carrive, P., Leung, P., Harris, J., & Paxinos, G. (1997). Conditioned fear to context is associated with increased Fos expression in the caudal ventrolateral region of the midbrain periaqueductal gray. Neuroscience, 78(1), 165–177. https://doi.org/10.1016/S0306-4522(97)83047-3

Chang, F. C. T. (1992). Modification of medullary respiratory-related discharge patterns by behaviors and states of arousal. Brain Research, 571(2), 281–292. https://doi.org/10.1016/0006-8993(92)90666-W

Chapuis, J., Cohen, Y., He, X., Zhang, Z., Jin, S., Xu, F., & Wilson, D. A. (2013). Lateral entorhinal modulation of piriform cortical activity and fine odor discrimination. Journal of Neuroscience, 33(33), 13449–13459. DOI: https://doi.org/10.1523/JNEUROSCI.1387-13.2013

Corfield, D. R., Murphy, K., & Guz, A. (1998). Does the motor cortical control of the diaphragm ‘bypass’ the brain stem respiratory centres in man?. Respiration Physiology, 114(2), 109–117. https://doi.org/10.1016/S0034-5687(98)00083-8

Dampney, R. A., Furlong, T. M., Horiuchi, J., & Iigaya, K. (2013). Role of dorsolateral periaqueductal grey in the coordinated regulation of cardiovascular and respiratory function. Autonomic Neuroscience, 175(1-2), 17–25. https://doi.org/10.1016/j.autneu.2012.12.008

Dampney, R. A. (2015). Central mechanisms regulating coordinated cardiovascular and respiratory function during stress and arousal. American Journal of Physiology-Regulatory, Integrative and Comparative Physiology, 309(5), R429–R443. https://doi.org/10.1152/ajpregu.00051.2015

Davis, P. J., Zhang, S. P., Winkworth, A., & Bandler, R. (1996). Neural control of vocalization: Respiratory and emotional influences. Journal of Voice, 10(1), 23–38. https://doi.org/10.1016/S0892-1997(96)80016-6

Del Negro, C. A., Funk, G. D., & Feldman, J. L. (2018). Breathing matters. Nature Reviews Neuroscience, 19, 351–367. https://doi.org/10.1038/s41583-018-0003-6

Depaulis, A., Bandler, R., & Vergnes, M. (1989). Characterization of pretentorial periaqueductal gray matter neurons mediating intraspecific defensive behaviors in the rat by microinjections of kainic acid. Brain research, 486(1), 121–132. https://doi.org/10.1016/0006-8993(89)91284-5

Deschênes, M., Moore, J., & Kleinfeld, D. (2012). Sniffing and whisking in rodents. Current Opinion in Neurobiology 22, 243–250. https://doi.org/10.1016/j.conb.2011.11.013

Dhingra, R. R., Furuya, W. I., Bautista, T. G., Dick, T. E., Galán, R. F., & Dutschmann, M. (2019a). Increasing local excitability of brainstem respiratory nuclei reveals a distributed network underlying respiratory motor pattern formation. Frontiers in Physiology, 10, 887. https://doi.org/10.3389/fphys.2019.00887

Dhingra, R. R., Furuya, W. I., Galán, R. F., & Dutschmann, M. (2019b). Excitation-inhibition balance regulates the patterning of spinal and cranial inspiratory motor outputs in rats in situ. Respiratory Physiology & Neurobiology, 266, 95–102. https://doi.org/10.1016/j.resp.2019.05.001

Dhingra, R. R., Dick, T. E., Furuya, W. I., Galán, R. F., & Dutschmann, M. (2020). Volumetric mapping of the functional neuroanatomy of the respiratory network in the perfused brainstem preparation of rats. The Journal of Physiology. https://doi.org/10.1113/JP279732.

Dick, T. E., Oku, Y., Romaniuk, J. R., & Cherniack, N. S. (1993). Interaction between central pattern generators for breathing and swallowing in the cat. The Journal of Physiology, 465(1), 715–730. https://doi.org/10.1113/jphysiol.1993.sp019702

Dillingham, C. M., Jankowski, M. M., Chandra, R., Frost, B. E., & O’Mara, S. M. (2017). The claustrum: considerations regarding its anatomy, functions and a programme for research. Brain and Neuroscience Advances, 1, 2398212817718962. https://doi.org/10.1177/2398212817718962.

Düsterhöft, F., Häusler, U., & Jürgens, U. (2004). Neuronal activity in the periaqueductal gray and bordering structures during vocal communication in the squirrel monkey. Neuroscience, 123(1), 53–60. https://doi.org/10.1016/j.neuroscience.2003.07.007

Dutschmann, M., & Paton, J. F. (2002). Inhibitory synaptic mechanisms regulating upper airway patency. Respiratory Physiology & Neurobiology, 131(1-2), 57–63. https://doi.org/10.1016/S1569-9048(02)00037-X

Dutschmann, M, & Herbert, H. (2006) The Kölliker-Fuse nucleus gates the postinspiratory phase of the respiratory cycle to control inspiratory off-switch and upper airway resistance in rat. European Journal of Neuroscience, 24, 1071–1084. https://doi.org/10.1111/j.1460-9568.2006.04981.x

Dutschmann, M., & Dick, T.E. (2012) Pontine mechanisms of respiratory control. Comprehensive Physiology, 2, 2443–2469. https://doi.org/10.1002/cphy.c100015

Dutschmann, M., Jones, S.E., Subramanian, H.H., Stanić, D., & Bautista, T.G. (2014) The physiological significance of postinspiration in respiratory control. In Progress in Brain Research, 212, 113–130. https://doi.org/10.1016/B978-0-444-63488-7.00007-0.

Dutschmann, M., Bautista, TG, Trevizan-Baú, P., Dhingra, R. R., Furuya, W. I. (2020). The pontine Kölliker-Fuse nucleus gates facial, hypoglossal, and vagal upper airway related motor activity. Respiratory Physiology & Neurobiology, to be submitted.

Farmer, D. G., Bautista, T. G., Jones, S. E., Stanić, D., & Dutschmann, M. (2014). The midbrain periaqueductal grey has no role in the generation of the respiratory motor pattern, but provides command function for the modulation of respiratory activity. Respiratory Physiology & Neurobiology, 204, 14–20. https://doi.org/10.1016/j.resp.2014.07.011

Faull, O. K., Subramanian, H. H., Ezra, M., & Pattinson, K. T. (2019). The midbrain periaqueductal gray as an integrative and interoceptive neural structure for breathing. Neuroscience & Biobehavioral Reviews, 98, 135–144. https://doi.org/10.1016/j.neubiorev.2018.12.020.

Feldman, J. L., & Del Negro, C. A. (2006). Looking for inspiration: new perspectives on respiratory rhythm. Nature Reviews Neuroscience, 7, 232–241. https://doi.org/10.1038/nrn1871

Finkelstein, D. I., Stanic, D., Parish, C. L., Tomas, D., Dickson, K., & Horne, M. K. (2000). Axonal sprouting following lesions of the rat substantia nigra. Neuroscience, 97, 99–112. https://doi.org/10.1016/s0306-4522(00)00009-9.

Fukushi, I., Yokota, S., & Okada, Y. (2019). The role of the hypothalamus in modulation of respiration. Respiratory Physiology & Neurobiology, 265, 172–179. https://doi.org/10.1016/j.resp.2018.07.003

Gandevia, S. C., & Rothwell, J. C. (1987a). Knowledge of motor commands and the recruitment of human motoneurons. Brain, 110(5), 1117–1130. https://doi.org/10.1093/brain/110.5.1117

Gandevia, S. C., & Rothwell, J. C. (1987b). Activation of the human diaphragm from the motor cortex. The Journal of Physiology, 384(1), 109–118. https://doi.org/10.1113/jphysiol.1987.sp016445

Geerling, J. C., Shin, J. W., Chimenti, P. C., & Loewy, A. D. (2010). Paraventricular hypothalamic nucleus: axonal projections to the brainstem. Journal of Comparative Neurology, 518(9), 1460–1499. https://doi.org/10.1002/cne.22283

Grady, F., Peltekian, L., Iverson, G., & Geerling, J. C. (2020). Direct Parabrachial–Cortical Connectivity. Cerebral Cortex. https://doi.org/10.1093/cercor/bhaa072.

Harper, R. M., Frysinger, R. C., Trelease, R. B., & Marks, J. D. (1984). State-dependent alteration of respiratory cycle timing by stimulation of the central nucleus of the amygdala. Brain Research, 306(1-2), 1–8. https://doi.org/10.1016/0006-8993(84)90350-0

Hage, S. R., & Nieder, A. (2016). Dual neural network model for the evolution of speech and language. Trends in neurosciences, 39(12), 813–829. https://doi.org/10.1016/j.tins.2016.10.006

Heck, D. H., McAfee, S. S., Liu, Y., Babajani-Feremi, A., Rezaie, R., Freeman, W. J., Wheless, W. J., Papanicolaou, A. C., Ruszinkó, M., Sokolov, Y., & Kozma, R. (2017). Breathing as a fundamental rhythm of brain function. Frontiers in Neural Circuits, 10, 115. https://doi.org/10.3389/fncir.2016.00115

Heck, D. H., Kozma, R., & Kay, L. M. (2019). The rhythm of memory: how breathing shapes memory function. Journal of Neurophysiology, 122(2), 563–571. https://doi.org/10.1152/jn.00200.2019

Hermann, D. M., Luppi, P. H., Peyron, C., Hinckel, P., & Jouvet, M. (1997). Afferent projections to the rat nuclei raphe magnus, raphe pallidus and reticularis gigantocellularis pars α demonstrated by iontophoretic application of choleratoxin (subunit b). Journal of Chemical Neuroanatomy, 13, 1–21. https://doi.org/10.1016/S0891-0618(97)00019-7

Hodges, M. R., & Richerson, G. B. (2010). The role of medullary serotonin (5-HT) neurons in respiratory control: contributions to eupneic ventilation, CO2 chemoreception, and thermoregulation. Journal of Applied Physiology, 108(5), 1425–1432. https://journals.physiology.org/doi/full/10.1152/japplphysiol.01270.2009

Holstege, G. (1989). Anatomical study of the final common pathway for vocalization in the cat. Journal of Comparative Neurology, 284(2), 242–252. https://doi.org/10.1002/cne.902840208

Holstege, G. (2014) The periaqueductal gray controls brainstem emotional motor systems including respiration. In Progress in Brain Research, 209, 379–405. https://doi.org/10.1016/B978-0-444-63274-6.00020-5

Holstege, G., & Subramanian, H. H. (2016). Two different motor systems are needed to generate human speech. Journal of Comparative Neurology, 524(8), 1558–1577. https://doi.org/10.1002/cne.23898

Holtman, J. R., Dick, T. E., & Berger, A. J. (1986). Involvement of serotonin in the excitation of phrenic motoneurons evoked by stimulation of the raphe obscurus. Journal of Neuroscience, 6(4), 1185–1193. https://doi.org/10.1523/JNEUROSCI.06-04-01185.1986

Homma, I., & Masaoka, Y. (2008). Breathing rhythms and emotions. Experimental Physiology, 93(9), 1011–1021. https://doi.org/10.1113/expphysiol.2008.042424

Jean, A. (2001). Brain stem control of swallowing: neuronal network and cellular mechanisms. Physiological Reviews, 81(2), 929–969. https://doi.org/10.1152/physrev.2001.81.2.929

Jürgens, U., & Richter, K. (1986). Glutamate-induced vocalization in the squirrel monkey. Brain Research, 373(1-2), 349–358. https://doi.org/10.1016/0006-8993(86)90349-5

Jürgens, U. (1994). The role of the periaqueductal grey in vocal behaviour. Behavioural Brain Research, 62(2), 107–117. https://doi.org/10.1016/0166-4328(94)90017-5

Jürgens, U. (2002). A study of the central control of vocalization using the squirrel monkey. Medical Engineering & Physics, 24(7-8), 473–477. https://doi.org/10.1016/S1350-4533(02)00051-6

Jürgens, U. (2009) The neural control of vocalization in mammals: a review. Journal of Voice, 23, 1–10. https://doi.org/10.1016/j.jvoice.2007.07.005

Larson, C. R. (1985). The midbrain periaqueductal gray: a brainstem structure involved in vocalization. Journal of Speech, Language, and Hearing Research, 28(2), 241–249. https://doi.org/10.1044/jshr.2802.241

Laplagne, D. A. (2018). Interplay between mammalian ultrasonic vocalizations and respiration. In Handbook of Behavioral Neuroscience (Vol. 25, pp. 61–70). Elsevier. https://doi.org/10.1016/B978-0-12-809600-0.00006-8

Leitner, F. C., Melzer, S., Lütcke, H., Pinna, R., Seeburg, P. H., Helmchen, F., & Monyer, H. (2016). Spatially segregated feedforward and feedback neurons support differential odor processing in the lateral entorhinal cortex. Nature neuroscience, 19(7), 935–944. https://doi.org/10.1038/nn.4303

Lindsey, B. G., Arata, A., Morris, K. F., Hernandez, Y. M., & Shannon, R. (1998). Medullary raphe neurones and baroreceptor modulation of the respiratory motor pattern in the cat. The Journal of Physiology, 512(3), 863–882. https://doi.org/10.1111/j.1469-7793.1998.863bd.x

Ludlow, C. L. (2015). Laryngeal reflexes: physiology, technique and clinical use. Journal of clinical neurophysiology, 32(4), 284-293. 10.1097/WNP.0000000000000187

Marchenko, V., Koizumi, H., Mosher, B., Koshiya, N., Tariq, M. F., Bezdudnaya, T. G., Zhang, R., Molkov, Y. I., Rybak, I. A., & Smith, J. C. (2016). Perturbations of respiratory rhythm and pattern by disrupting synaptic inhibition within pre-Bötzinger and Bötzinger complexes. ENeuro, 3(2). 10.1523/ENEURO.0011-16.2016

Mazzone, S. B., Cole, L. J., Ando, A., Egan, G. F., & Farrell, M. J. (2011). Investigation of the neural control of cough and cough suppression in humans using functional brain imaging. Journal of Neuroscience, 31(8), 2948–2958. https://doi.org/10.1523/JNEUROSCI.4597-10.2011

McElvain, L. E., Friedman, B., Karten, H. J., Svoboda, K., Wang F., Deschênes M., Kleinfeld, D. (2018). Circuits in the rodent brainstem that control whisking in concert with other orofacial motor actions. Neuroscience, 368, 152–170. https://doi.org/10.1016/j.neuroscience.2017.08.034

Moore, J. D., Deschênes, M., Furuta, T., Huber, D., Smear, M. C., Demers, M., & Kleinfeld, D. (2013). Hierarchy of orofacial rhythms revealed through whisking and breathing. Nature, 497(7448), 205–210. https://doi.org/10.1038/nature12076

Moore, J. D., Kleinfeld, D., & Wang, F. (2014). How the brainstem controls orofacial behaviors comprised of rhythmic actions. Trends in Neurosciences, 37, 370–380. https://doi.org/10.1016/j.tins.2014.05.001

Mörschel, M., & Dutschmann, M. (2009). Pontine respiratory activity involved in inspiratory/expiratory phase transition. Philosophical Transactions of the Royal Society B: Biological Sciences, 364(1529), 2517–2526. https://doi.org/10.1098/rstb.2009.0074

Nonaka, S., Sakamoto, T., Katada, A., & Unno, T. (1999). Brain stem neural mechanisms for vocalization in decerebrate cats. Annals of Otology, Rhinology & Laryngology, 108(7_suppl), 15–24. https://doi.org/10.1177/00034894991080S703

Onimaru, H., & Homma, I. (2007). Spontaneous oscillatory burst activity in the piriform–amygdala region and its relation to in vitro respiratory activity in newborn rats. Neuroscience, 144(1), 387–394. https://doi.org/10.1016/j.neuroscience.2006.09.033

Orem, J., & Netick, A. (1986). Behavioral control of breathing in the cat. Brain research, 366(1-2), 238–253. https://doi.org/10.1016/0006-8993(86)91301-6

Paxinos, G., & Watson, C. (2007). The Rat Brain in Stereotaxic Coordinates: Academic Press.

Pérez de los Cobos Pallares, F., Bautista, T. G., Stanić, D., Egger, V., & Dutschmann, M. (2016). Brainstem-mediated sniffing and respiratory modulation during odor stimulation. Respiratory Physiology & Neurobiology, 233, 17–24. https://doi.org/10.1016/j.resp.2016.07.008

Petersen, C. C. (2019). Sensorimotor processing in the rodent barrel cortex. Nature Reviews Neuroscience, 20(9), 533–546. https://doi.org/10.1038/s41583-019-0200-y

Peyron, C., Tighe, D. K., Van Den Pol, A. N., De Lecea, L., Heller, H. C., Sutcliffe, J. G., & Kilduff, T. S. (1998). Neurons containing hypocretin (orexin) project to multiple neuronal systems. Journal of Neuroscience, 18(23), 9996–10015. https://doi.org/10.1523/JNEUROSCI.18-23-09996.1998

Pouget, P., Allard, E., Poitou, T., Raux, M., Wattiez, N., & Similowski, T. (2018). Slower is higher: Threshold modulation of cortical activity in voluntary control of breathing initiation. Frontiers in Neuroscience, 12, 663. https://doi.org/10.3389/fnins.2018.00663

Richter, D. W., Manzke, T., Wilken, B., & Ponimaskin, E. (2003). Serotonin receptors: guardians of stable breathing. Trends in Molecular Medicine, 9(12), 542–548. https://doi.org/10.1016/j.molmed.2003.10.010

Rikard-Bell, G. C., Bystrzycka, E. K., & Nail, B. S. (1985). Cells of origin of corticospinal projections to phrenic and thoracic respiratory motoneurones in the cat as shown by retrograde transport of HRP. Brain Research Bulletin, 14(1), 39–47. https://doi.org/10.1016/0361-9230(85)90175-3

Ruggiero, D. A., Mraovitch, S., Granata, A. R., Anwar, M., & Reis, D. J. (1987). A role of insular cortex in cardiovascular function. Journal of Comparative Neurology, 257(2), 189–207. https://doi.org/10.1002/cne.902570206

Semba, K., & Egger, M. D. (1986). The facial “motor” nerve of the rat: control of vibrissal movement and examination of motor and sensory components. Journal of Comparative Neurology, 247(2), 144–158. https://doi.org/10.1002/cne.902470203

Saper, C. B. (1982). Convergence of autonomic and limbic connections in the insular cortex of the rat. Journal of Comparative Neurology, 210(2), 163–173. https://doi.org/10.1002/cne.902100207

Sato, F., Akhter, F., Haque, T., Kato, T., Takeda, R., Nagase, Y., Sessle, B. J., & Yoshida, A. (2013). Projections from the insular cortex to pain-receptive trigeminal caudal subnucleus (medullary dorsal horn) and other lower brainstem areas in rats. Neuroscience, 233, 9–27. https://doi.org/10.1016/j.neuroscience.2012.12.024

Sherwood, C. C., Hof, P. R., Holloway, R. L., Semendeferi, K., Gannon, P. J., Frahm, H. D., & Zilles, K. (2005). Evolution of the brainstem orofacial motor system in primates: a comparative study of trigeminal, facial, and hypoglossal nuclei. Journal of Human Evolution, 48, 45–84. https://doi.org/10.1016/j.jhevol.2004.10.003

Smith, J. C., Ellenberger, H. H., Ballanyi, K., Richter, D. W., & Feldman, J. L. (1991). Pre-Botzinger complex: a brainstem region that may generate respiratory rhythm in mammals. Science, 254, 726-729. 10.1126/science.1683005

Smith, J. C., Abdala, A. P. L., Koizumi, H., Rybak, I. A., & Paton, J. F. (2007). Spatial and functional architecture of the mammalian brain stem respiratory network: a hierarchy of three oscillatory mechanisms. Journal of Neurophysiology. https://doi.org/10.1152/jn.00985.2007

Smith, J. C., Abdala, A. P., Borgmann, A., Rybak, I. A., & Paton, J. F. (2013). Brainstem respiratory networks: building blocks and microcircuits. Trends in neurosciences, 36(3), 152–162. https://doi.org/10.1016/j.tins.2012.11.004

Stanic, D., Finkelstein, D. I., Bourke, D. W., Drago, J., & Horne, M. K. (2003). Timecourse of striatal re-innervation following lesions of the dopaminergic SNpc neurons of the rat. European Journal of Neuroscience, 18, 1175–88. https://doi.org/10.1046/j.1460-9568.2003.02800.x

Steinmetz, N. A., Zatka-Haas, P., Carandini, M., & Harris, K. D. (2019). Distributed coding of choice, action and engagement across the mouse brain. Nature, 576(7786), 266–273. https://doi.org/10.1038/s41586-019-1787-x

Subramanian, H. H., Balnave, R. J., & Holstege, G. (2008a). The midbrain periaqueductal gray control of respiration. Journal of Neuroscience, 28(47), 12274–12283. https://doi.org/10.1523/JNEUROSCI.4168-08.2008

Subramanian, H., Huang, Z. G., & Balnave, R. (2008b). Responses of brainstem respiratory neurons to activation of midbrain periaqueductal gray in the rat. In Integration in Respiratory Control (pp. 377–381). Springer, New York, NY.

Subramanian, H. H., & Holstege, G. (2009). The nucleus retroambiguus control of respiration. Journal of Neuroscience, 29(12), 3824–3832. https://doi.org/10.1523/JNEUROSCI.0607-09.2009

Subramanian, H. H., & Holstege, G. (2014). The midbrain periaqueductal gray changes the eupneic respiratory rhythm into a breathing pattern necessary for survival of the individual and of the species. In Progress in Brain Research, 212, 351–384. https://doi.org/10.1016/B978-0-444-63488-7.00017-3

Trevizan-Baú, P, Furuya WI, Mazzone SB, Stanić, D, Dhingra RR, Dutschmann, M. (2020). Reciprocal connectivity of the lateral and ventrolateral columns of the midbrain periaqueductal gray with core nuclei of ponto-medullary respiratory network in rat. Brain Research, to be submitted.

Verdejo-Garcia, A., Clark, L., & Dunn, B. D. (2012). The role of interoception in addiction: a critical review. Neuroscience & Biobehavioral Reviews, 36(8), 1857–1869. https://doi.org/10.1016/j.neubiorev.2012.05.007

Wang, W., Fung, M. L., & St John, W. M. (1993). Pontile regulation of ventilatory activity in the adult rat. Journal of Applied Physiology, 74(6), 2801–2811. https://doi.org/10.1152/jappl.1993.74.6.2801

Woolsey, T. A., & Van der Loos, H. (1970). The structural organization of layer IV in the somatosensory region (SI) of mouse cerebral cortex: the description of a cortical field composed of discrete cytoarchitectonic units. Brain research, 17(2), 205–242. https://doi.org/10.1016/0006-8993(70)90079-X

Yasui, Y., Breder, C. D., Safer, C. B., & Cechetto, D. F. (1991). Autonomic responses and efferent pathways from the insular cortex in the rat. Journal of Comparative Neurology, 303(3), 355–374. https://doi.org/10.1002/cne.903030303

Zald, D. H., & Pardo, J. V. (1999). The functional neuroanatomy of voluntary swallowing. Annals of Neurology: Official Journal of the American Neurological Association and the Child Neurology Society, 46(3), 281–286. https://doi.org/10.1002/1531-8249(199909)46:3%3C281::AID-ANA2%3E3.0.CO;2-L

Zhang, S. P., Davis, P. J., Bandler, R., & Carrive, P. (1994). Brain stem integration of vocalization: role of the midbrain periaqueductal gray. Journal of Neurophysiology, 72(3), 1337–1356. https://doi.org/10.1152/jn.1994.72.3.1337

Zhang, S. P., Bandler, R., & Davis, P. J. (1995). Brain stem integration of vocalization: role of the nucleus retroambigualis. Journal of Neurophysiology, 74(6), 2500–2512. https://doi.org/10.1152/jn.1995.74.6.2500

